# Chromatin modifiers KMT2D, BAF, and p300 are required for *de novo* binding of transcription factors on enhancers

**DOI:** 10.64898/2026.01.29.702555

**Authors:** Hieu T. Van, Young-Kwon Park, Chengyu Liu, Shamima Islam, Stefania Dell’Orso, Weiqun Peng, Vittorio Sartorelli, Ji-Eun Lee, Kai Ge

## Abstract

Transcription factors (TFs) bind to enhancers and recruit H3K4me1 methyltransferase KMT2D, chromatin remodeler BAF, and H3K27 acetyltransferase p300 to activate transcription. However, the role of chromatin modifiers in regulating *de novo* binding of TFs on enhancers remains unclear. Using a robust nuclear translocation system, we show that the muscle lineage-determining TF MyoD binds to chromatin pervasively within one hour, with half of induced MyoD binding sites co-occupied by KMT2D, BAF, and p300. On the majority of these MyoD^+^ enhancers, acute depletion of KMT2D or short-term inhibition of BAF or p300 enzymatic activity markedly reduces *de novo* binding of MyoD as well as that of KMT2D, BAF, and p300. On enhancers with intact MyoD binding despite these perturbations, we observe a cooperative recruitment among chromatin modifiers. Similar interdependent relationships are observed between the signal-dependent TF Glucocorticoid Receptor and KMT2D, BAF, and p300. Together, our findings show that chromatin modifiers are not only downstream effectors but also required for *de novo* binding of TFs on enhancers, refining a model of enhancer establishment as a process governed by functional cooperation rather than a strict hierarchy.

**Bullet points:**

- Acute KMT2D depletion disrupts *de novo* binding of MyoD, BAF, and p300 on enhancers.
- BAF and p300 enzymatic activities are required for *de novo* binding of MyoD, BAF, KMT2D, and p300 on enhancers.
- Cooperative binding of KMT2D, BAF, and p300 on MyoD^+^ enhancers.
- GR displays interdependencies with chromatin modifiers KMT2D, BAF, and p300 on enhancers.

## Introduction

Cell differentiation, the process by which precursor cells give rise to specialized cell types, is crucial for the development of multicellular organisms. Cell differentiation is determined by cell-type specific gene expression, which is precisely controlled by transcriptional enhancers. Enhancers are DNA elements enriched for binding motifs of lineage-determining transcription factors (LDTFs). Upon binding to enhancers, LDTFs recruit other TFs, coactivators, and chromatin modifiers, priming these regulatory regions for activation and facilitating RNA Polymerase II recruitment to drive the expression of cell-type specific genes [1]. Enhancers are often characterized by the presence of histone H3K4 mono-methylation (H3K4me1) in their primed states, or both H3K4me1 and H3K27 acetylation (H3K27ac) in their active states [1].

Key chromatin modifiers, including KMT2C/D, CBP/p300, and the BAF complex, play important roles in enhancer regulation. KMT2C/D (also known as MLL3/4), are major mammalian H3K4me1 methyltransferase enriched on enhancers [2]. KMT2C/D are critical for *de novo* enhancer activation during cell differentiation and cell fate transitions, with KMT2D holding a more prominent role [2, 3]. The histone acetyltransferase p300 (also known as EP300/KAT3B) and its homolog CBP (also known as CREBBP/KAT3A) catalyze H3K27ac [4] and mark cell-type specific enhancers in mammalian cells [5].

Beside acetylating histones, CBP/p300 contribute to transcription by acetylating non-histone proteins, scaffolding transcriptional machineries, and promoting enhancer-promoter communication [6]. The BAF complex, one of the three SWI/SNF chromatin remodelers, facilitates chromatin opening by mobilizing and evicting nucleosomes through its enzymatic subunit BRG1 (also known as SMARCA4) [7]. BAF colocalizes with LDTFs on enhancers and promotes enhancer activation during cell fate transitions [8]. Consistent with their critical roles in enhancer activation, KMT2C/D, CBP/p300, and BAF are generally required for cell differentiation and animal development [9, 10]. Mutations in the encoding genes of these chromatin modifiers are found to drive many types of cancers [11].

Our previous studies, using various cell differentiation models including adipogenesis, myogenesis, and mouse embryonic stem cell (ESC) differentiation, have identified functional relationships among KMT2D, p300, and BAF on enhancers. Specifically, loss of KMT2D disrupts p300 recruitment and impairs *de novo* enhancer activation [2, 3, 12], suggesting that p300 functions downstream of KMT2D. KMT2D also physically associates with BAF, and loss of BAF’s ATPase subunit BRG1 blocks activation of C/EBPβ-bound adipogenic enhancers. BAF and KMT2D reciprocally regulate each other’s binding on enhancers [8]. These findings suggest a stepwise model of enhancer activation: (1) LDTFs bind to enhancer regions; (2) BAF and KMT2D are recruited; and (3) p300 is recruited to promote enhancer activation [13]. However, these insights are largely derived from stable gene knockout approaches, which allow for compensatory or indirect effects that might obscure the primary functions of these chromatin modifiers. For example, BRG1 has been reported to act as both a repressor and an activator of transcription in knockout studies, whereas acute depletion of BRG1 proteins indicates that it primarily functions as a transcriptional activator [14]. Additionally, previous studies used models in which LDTFs are stably expressed, resulting in a pre-established enhancer landscape, which limits the analysis of *de novo* binding events.

MyoD is the founding member of the myogenic TF family [15] and plays a crucial role in coordinating transcriptional programs and promoting myogenesis in most cell types [16]. Chromatin modifiers BAF, KMT2D, and p300 are known to be important for MyoD-mediated enhancer activation, gene induction, and myogenesis [2, 17–19]. To overcome limitations of previous studies and to investigate the early dynamics and functional relationships of LDTFs and chromatin modifiers on enhancers, we used a robust MyoD nuclear translocation system, coupled with short-term depletion of KMT2D or chemical inhibition of BAF or p300 enzymatic activity. We found that within one hour (1h) of its nuclear translocation, MyoD colocalizes with BAF, KMT2D, and p300 on approximately 50% of enhancers. Acute perturbation of any of these chromatin modifiers disrupts *de novo* binding of MyoD on most MyoD^+^ enhancers. On enhancers where MyoD binding remained intact, acute perturbation of one chromatin modifier markedly reduced the binding of the others, suggesting that KMT2D, BAF, and p300 regulate one another on LDTF-bound enhancers. In addition, we used an endogenous nuclear translocation system of the signal-dependent TF (SDTF) Glucocorticoid Receptor (GR) and validated the dynamic and interdependent relationships between TFs and chromatin modifiers on enhancers.

## Results

### Dynamic binding of nuclear translocated MyoD during myogenesis

To overcome the limitations inherent to stably expressed LDTF models and to elucidate the early interplay among LDTFs and chromatin modifiers on enhancers, we utilized a 4-Hydroxytamoxifen (4-OHT)-inducible MyoD-ER nuclear translocation system [20]. We generated a retroviral vector carrying MyoD-ER with three tandem T7 tags at the C-terminus (Fig. 1A) and expressed MyoD-ER-T7 in preadipocytes, where endogenous *MyoD* expression was undetectable (Fig. 1B). 4-OHT treatment markedly increased nuclear MyoD levels within 1h (Fig. 1C). Similar to constitutively expressed MyoD [2], nuclear translocated MyoD-ER-T7 could induce the *trans*-differentiation of preadipocytes into myocytes and the associated expression of myogenic marker genes *Myogenin (Myog)* and *Creatine kinase (Ckm)* (Fig. 1-S1A-B). We hereafter refer to nuclear translocated MyoD-ER-T7 as MyoD.

**Fig 1.**
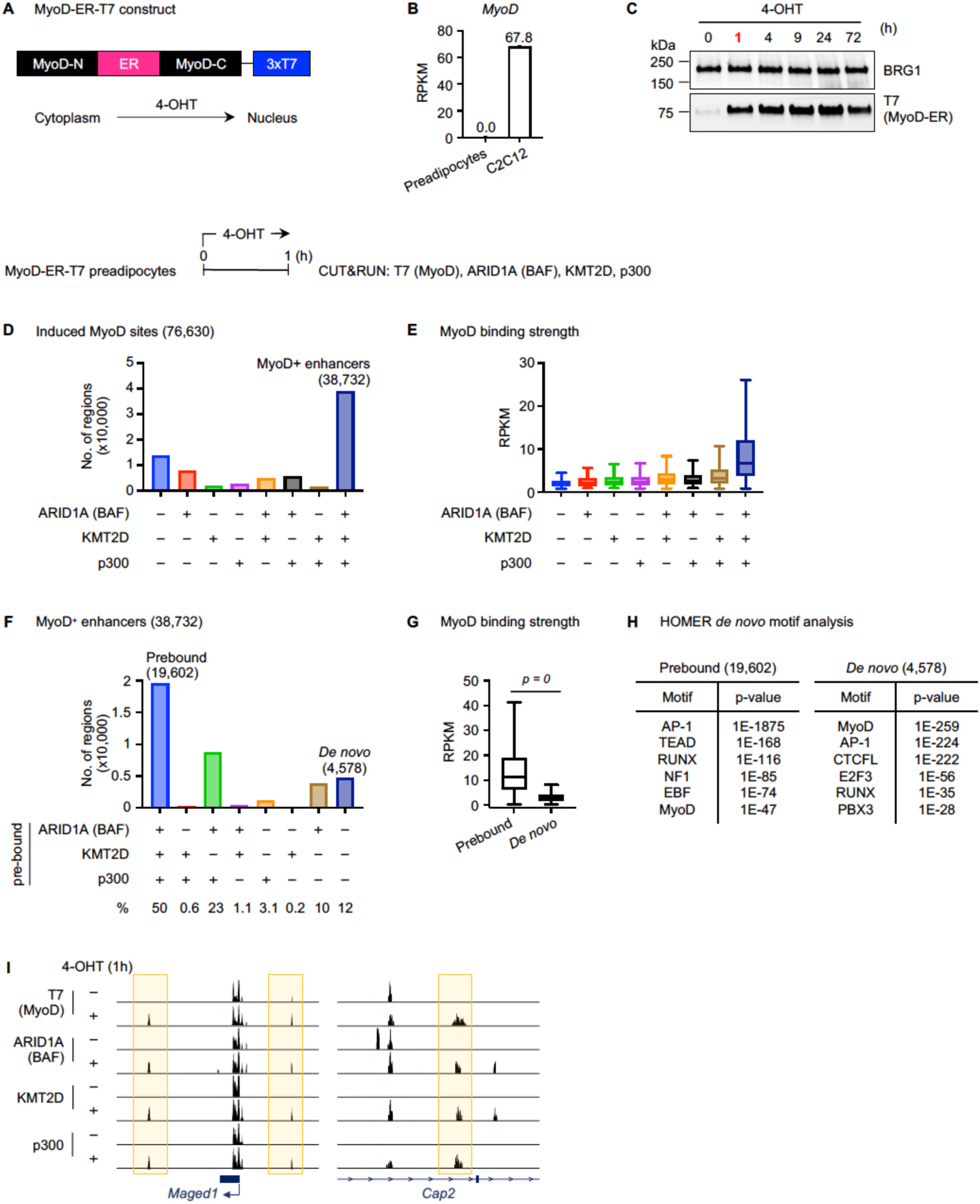
Acutely translocated MyoD colocalizes with chromatin modifiers BAF, KMT2D, and p300 on enhancers. **(A)** Schematic of the 4-OHT-inducible MyoD-ER system. **(B)** *MyoD* expression in preadipocytes and C2C12 myoblasts was determined using RNA-Seq (n = 1). RPKM values indicate gene expression levels. **(C)** Western blot (WB) analysis of nuclear extracts from preadipocytes expressing MyoD-ER-T7 and treated with 4-OHT. Antibodies used were indicated on the right. BRG1 was used as a loading control. **(D-I)** MyoD-ER-T7 expressing preadipocytes were treated with 4-OHT for 1 hour (h), followed by CUT&RUN analysis. **(D)** Bar chart showing ARID1A (an exclusive subunit of BAF), KMT2D, and p300 binding status on induced MyoD sites. **(E)** Box plots displaying the normalized MyoD read counts in subgroups defined in (D). **(F)** Bar chart showing ARID1A (BAF), KMT2D, and p300 binding on 38,732 MyoD^+^ enhancers defined in (D) prior to 4-OHT treatment. **(G-H)** Box plots showing the normalized MyoD read counts **(G)** and HOMER *de* novo motif analysis **(H)** on BAF-KMT2D-p300 prebound or *de novo* sites defined in (F). Statistical significance was determined using a two-sided, unpaired Mann Whitney test. **(I)** Genome browser view of MyoD binding sites around *Maged1* and *Cap2* loci.

To characterize the binding dynamics of nuclear translocated MyoD during myogenic *trans*-differentiation, we performed time-course analysis using Cleavage Under Targets and Release Using Nuclease (CUT&RUN) with a T7 antibody. We identified induced MyoD binding sites upon 4-OHT treatment at each time point, where the signal intensity was >2-fold higher than that at 0h. Motif analysis confirmed MyoD as the most significantly enriched motif within induced MyoD binding sites across all time points (Fig. 1-S1C). On the *Myog* locus, MyoD binding was significant only after 24h of 4-OHT treatment, whereas on the surrounding regions, MyoD binding appeared as early as at 1h (Fig. 1-S1D). We then identified 36,425 induced MyoD binding sites overlapped with H3K4me1^+^ or H3K27ac^+^ regions (Fig. 1-S1E). Unbiased K-means clustering analysis revealed six distinct clusters A-F, each with a unique MyoD binding pattern (Fig. 1-S1F). MyoD binding near muscle cell differentiation and development genes peaked at 24h or later in clusters D and E. In contrast, early MyoD binding sites in clusters A, B, and C were associated with genes involved in more general cellular processes such as cell-substrate adhesion (Fig. 1-S1G). These results suggest that MyoD targets distinct enhancers in a temporally regulated manner during myogenic reprogramming of preadipocytes.

### Acutely translocated MyoD colocalizes with chromatin modifiers on enhancers

Because the majority of MyoD-ER is translocated into the nucleus within 1h of 4-OHT treatment (Fig. 1C), we focused on induced MyoD sites at 1h to investigate the early binding of MyoD and chromatin modifiers. We examined the genomic enrichment of KMT2D, BAF, and p300 on MyoD binding sites, using CUT&RUN with antibodies against T7 tag (for MyoD), ARID1A (a representative and exclusive subunit of BAF), KMT2D, and p300. Among the 76,630 induced MyoD binding sites, ∼51% (38,732 sites) were co-occupied by all three chromatin modifiers KMT2D, BAF, and p300 (Fig. 1D). The strength of MyoD binding on these 38,732 sites was significantly higher than on any other subset of sites (Fig. 1E). Because BAF, KMT2D, and p300 are well-established enhancer regulators, we called this group as MyoD⁺ enhancers. For subsequent analysis, we focused on these 38,732 MyoD^+^ enhancers.

Prior to MyoD recruitment, the majority of the 38,732 MyoD^+^ enhancers were prebound by at least one of the three chromatin modifiers, with 50% (19,602 sites) prebound by all three. In contrast, approximately 12% (4,578 sites) showed *de novo* binding of KMT2D, BAF, and p300 upon MyoD recruitment (Fig. 1F). MyoD binding was significantly stronger on BAF-KMT2D-p300 prebound sites compared to the *de novo* sites (Fig. 1G). Although the MyoD motif was enriched in both groups of sites, it was the most significantly enriched motif on the *de novo* sites, whereas the AP-1 motif ranked highest on prebound sites (Fig. 1H-I).

### Acute KMT2D depletion disrupts *de novo* binding of MyoD, BAF, and p300 on enhancers

To investigate the roles of KMT2D in regulating MyoD, BAF, and p300 binding on enhancers, we utilized the auxin-inducible degron 2 (AID2) system to acutely deplete KMT2D, the largest nuclear protein in mammalian cells [9, 21]. For this purpose, *Kmt2d* ^AID/AID^ mice expressing AID-tagged KMT2D protein were generated using CRISPR-mediated insertion of an AID tag to the N-terminus of *Kmt2d* alleles (Fig. 2A; Fig. 2-S1A). We then crossed *Kmt2d* ^AID/AID^ mice with *Kmt2c* ^f/f^; *Cre-ER* mice [22] to obtain primary *Kmt2c* ^f/f^; *Kmt2d* ^AID/AID^; *Cre-ER* preadipocytes (Fig. 2-S1B). After immortalization, cells were treated with 4-OHT to permanently delete the *Kmt2c* gene by Cre recombinase to prevent the potential functional compensation by KMT2C [2]. The resulting *Kmt2c* ^-/-^; *Kmt2d* ^AID/AID^ cells were infected with retroviruses expressing F74G-mutant *Oryza sativa* TIR1 (OsTIR1-F74G) and MyoD-ER-T7 (Fig. 2-S1C-D). Following 5Ph-IAA treatment, KMT2D was depleted within 2h, which was also confirmed by the loss of UTX, whose stability depends on KMT2D (Fig. 2-S1E) [23]. We refer to these cells as *Kmt2d* ^AID/AID^; MyoD-ER-T7.

**Fig 2.**
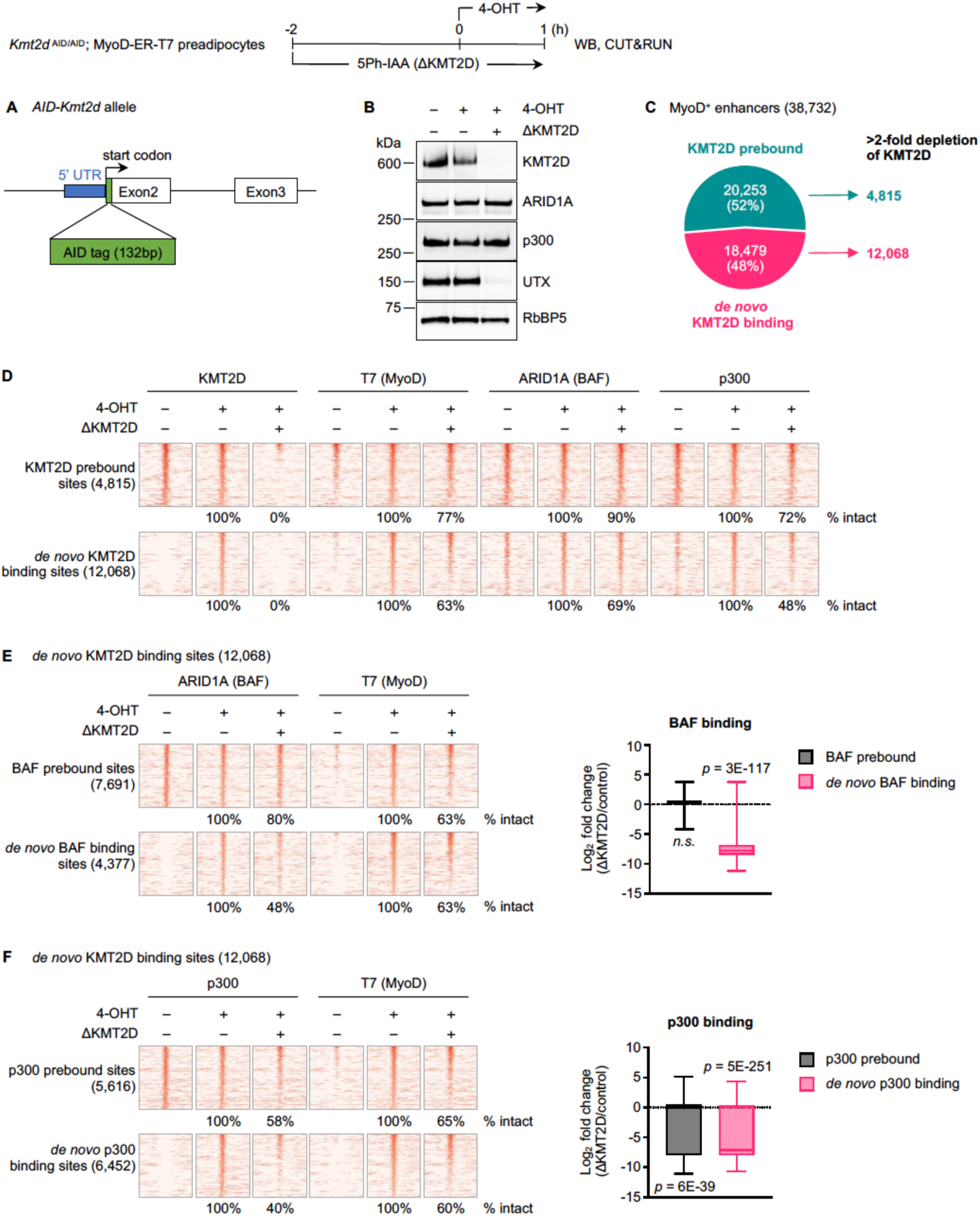
Acute KMT2D depletion disrupts *de novo* binding of MyoD, BAF, and p300 on enhancers. **(A)** Schematic for generating the knockin allele encoding AID-tagged KMT2D. **(B-F)** *Kmt2d* ^AID/AID^; MyoD-ER-T7 preadipocytes were pretreated with 5Ph-IAA (ΔKMT2D) for 2h, and then 4-OHT was added for 1h to induce MyoD nuclear translocation. Cells were harvested for WB and CUT&RUN analysis. **(B)** WB of nuclear extracts for KMT2D, ARID1A (BAF), p300, and UTX. Antibodies used were indicated on the right. RbBP5 was the loading control. **(C)** Pie chart illustrating KMT2D binding status on 38,732 MyoD^+^ enhancers. **(D)** Heat maps for CUT&RUN of KMT2D, T7 (MyoD), ARID1A (BAF), and p300 on KMT2D prebound or *de novo* KMT2D binding sites with >2-fold depletion of KMT2D as defined in (C). **(E-F)** Heat maps (*left panel)* for CUT&RUN data on 12,068 MyoD^+^ enhancers with *de novo* KMT2D binding, further categorized based on BAF binding **(E)** or p300 binding **(F)** before and after 4-OHT treatment. All heat maps spanned ± 3kb around MyoD binding sites, and sites were ranked by the intensity of MyoD (T7) in the 4OHT-treated control. Box plots (*right panel*) showing fold changes of BAF binding intensity **(E)** or p300 binding intensity **(F)** between KMT2D-depleted (ΔKMT2D) and control samples. Statistical significance was determined using a one-sided Wilcoxon signed-rank test.

*Kmt2d* ^AID/AID^; MyoD-ER-T7 cells were pretreated with 5Ph-IAA for 2h, and then 4-OHT was added for 1h to induce MyoD nuclear translocation. As shown in Fig. 2B, 3h of 5Ph-IAA treatment significantly depleted KMT2D and UTX without much effect on ARID1A and p300 protein levels. To examine how acute loss of KMT2D affects MyoD, BAF, and p300 binding, we performed CUT&RUN. On the 38,732 MyoD^+^ enhancers (Fig. 1D), we observed that 52% (20,253 sites) were prebound by KMT2D, while the remaining 48% (18,479 sites) showed *de novo* KMT2D binding. We focused on 4,815 KMT2D prebound sites and 12,068 *de novo* KMT2D binding sites where KMT2D signal intensity was reduced >2-fold upon depletion (Fig. 2C). On KMT2D prebound sites, the loss of KMT2D did not affect MyoD and p300 binding on >70% of the sites, and 90% of BAF binding sites were also intact. However, KMT2D depletion had a more significant impact on *de novo* KMT2D binding sites. 37% of these sites lost MyoD binding, whereas 31% and 52% showed reduced BAF and p300 binding, respectively (Fig. 2D).

The 12,068 *de novo* KMT2D binding sites were further categorized according to BAF or p300 binding status prior to MyoD translocation, as we observed that many of these sites were pre-marked by either of these factors. On BAF- or p300-prebound sites, KMT2D depletion reduced BAF or p300 binding on 20% or 42% of them, respectively. On sites with *de novo* BAF or p300 binding, however, the loss of KMT2D reduced BAF or p300 binding on 52% or 60% of them, respectively (Fig. 2E-F). Together, these results indicate that KMT2D is important for *de novo* binding of MyoD, BAF, and p300 on enhancers.

### BAF activity is required for *de novo* binding of MyoD and chromatin modifiers on enhancers

Next, we addressed the role of BAF in modulating the early binding of MyoD and chromatin modifiers on enhancers. We utilized BRM014 (BRG1i), a catalytic inhibitor of the BAF enzymatic subunit BRG1 [24, 25]. We pretreated *Kmt2d* ^AID/AID^; MyoD-ER-T7 cells with BRG1i for 1h and then induced MyoD nuclear translocation with 4-OHT for 1h. BRG1i treatment did not alter the protein levels of BRG1, ARID1A, KMT2D, or p300 (Fig. 3A).

**Fig 3.**
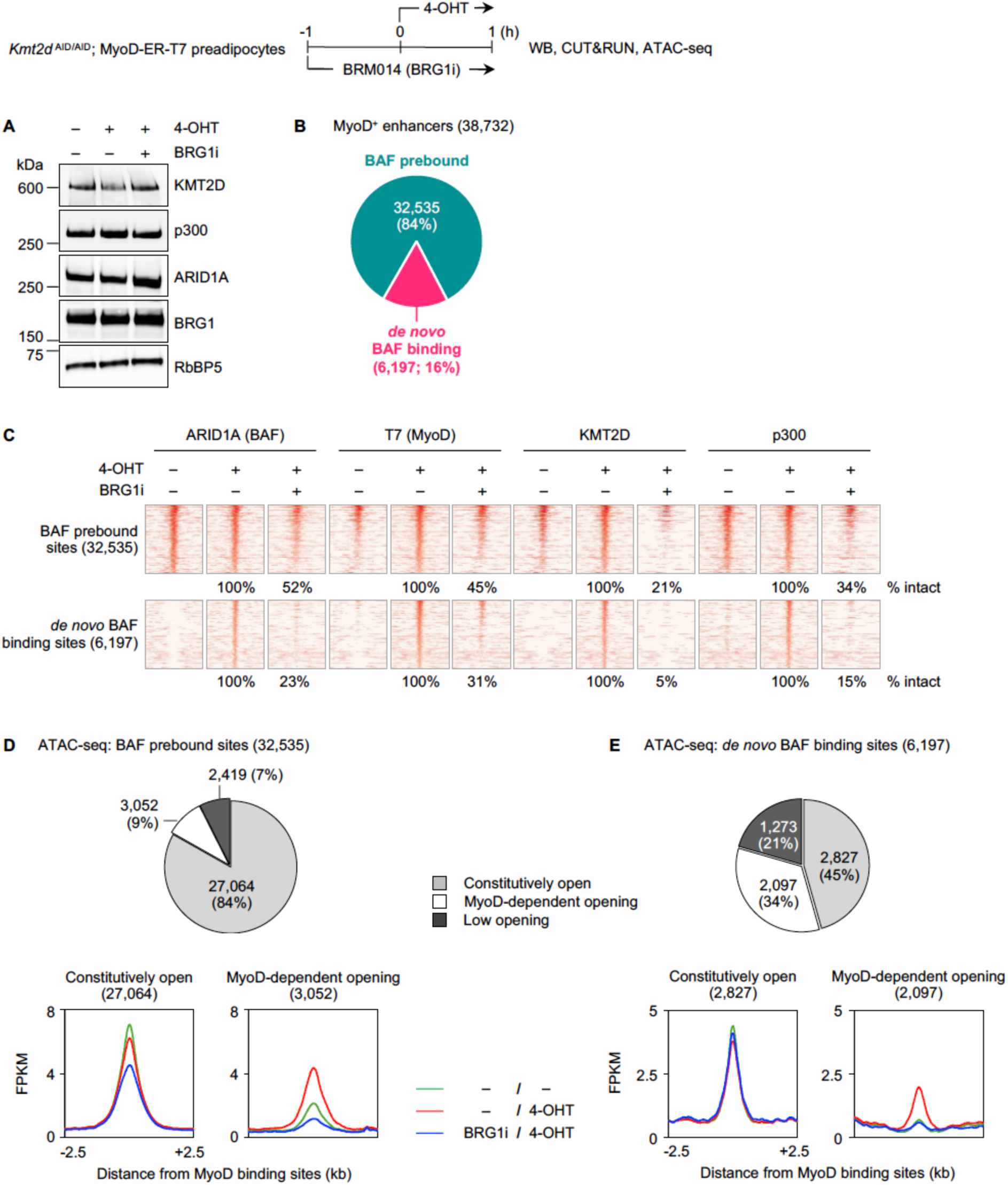
Inhibition of BAF activity markedly reduces *de novo* binding of MyoD, KMT2D, BAF, and p300 on enhancers. *Kmt2d* ^AID/AID^; MyoD-ER-T7 preadipocytes were pretreated with BRG1 inhibitor BRM014 (BRG1i) for 1h, and then 4-OHT was added for 1h to induce MyoD nuclear translocation. Cells were harvested for WB, CUT&RUN, and ATAC-seq. **(A)** WB of nuclear extracts for KMT2D, p300, and BAF subunits BRG1 and ARID1A. **(B)** Pie chart illustrating BAF binding status on 38,732 MyoD^+^ enhancers. **(C)** Heat maps for CUT&RUN of ARID1A (BAF), T7 (MyoD), KMT2D, and p300 on BAF pre-bound and *de novo* BAF binding sites. **(D-E)** Chromatin accessibility determined by ATAC-seq signals on MyoD^+^ enhancers. Chromatin accessibility status on BAF prebound sites **(D)** or *de novo* BAF binding sites **(E)** is shown in pie charts (*upper panels*). Average profiles of normalized ATAC-seq reads on constitutively open and MyoD-dependent opening sites are shown in *lower panels*.

We performed CUT&RUN to assess the effects of BRG1 inhibition on MyoD, BAF, KMT2D, and p300 chromatin binding. On the 38,732 MyoD^+^ enhancers (Fig. 1A), the vast majority (32,535) were prebound by BAF, while 16% (6,197) showed *de novo* BAF binding (Fig. 3B). On BAF prebound sites, BRG1i treatment led to a >2-fold decrease in MyoD binding on 55% of them. Binding of all chromatin modifiers was also broadly reduced. BAF, KMT2D, and p300 binding showed >2-fold reduction on 48%, 79%, and 66% of the prebound sites, respectively (Fig. 3C). The impact was even more pronounced on *de novo* BAF binding sites, where the loss of BRG1 activity diminished MyoD and BAF binding on 69% and 77% of sites and nearly abolished KMT2D and p300 binding on 95% and 85% of them, respectively (Fig. 3C).

To investigate chromatin accessibility on MyoD^+^ enhancers, we performed ATAC-seq and compared the signals before and after MyoD nuclear translocation. We identified three groups with distinct characteristics: constitutively open, MyoD-dependent opening, and low opening. Among 32,535 BAF prebound enhancers, the majority (84%; 27,064 sites) were constitutively open, while 9% (3,052 sites) and 7% (2,419 sites) belonged to the MyoD-dependent opening or low opening group, respectively (Fig. 3D, *upper panel*). In contrast, a significant number (34%; 2,097 sites) of *de novo* BAF binding sites showed MyoD-dependent opening, while 45% and 21% were constitutively or lowly open, respectively (Fig. 3E, *upper panel*). We then examined changes in chromatin accessibility upon BRG1i treatment. On BAF prebound enhancers, inhibition of BRG1 activity led to a more drastic decrease in chromatin accessibility on 3,052 MyoD-dependent opening sites, compared to that on the 27,064 constitutively open sites (Fig. 3D, *lower panel*). On *de novo* BAF binding sites, BRG1 inhibition did not alter chromatin accessibility on constitutively open sites, but completely abolished MyoD-mediated chromatin opening (Fig. 3E, *lower panel*). These results suggest that MyoD-dependent chromatin opening depends on BRG1 enzymatic activity. Our data also indicate that BRG1 activity is required for *de novo* binding of MyoD, BAF complex itself, and other chromatin modifiers KMT2D and p300 on enhancers.

### *De novo* binding of MyoD and chromatin modifiers depends on p300 acetyltransferase activity

To understand how p300 acetyltransferase activity influences MyoD and chromatin modifier binding on enhancers, we used a p300 catalytic inhibitor A485 (p300i) [26]. *Kmt2d* ^AID/AID^; MyoD-ER-T7 cells were pretreated with p300i for 1h, followed by 4-OHT treatment to induce MyoD nuclear translocation for 1h. p300i treatment did not affect protein levels of p300, KMT2D, and ARID1A, but significantly reduced global H3K27ac levels (Fig. 4A).

**Fig 4.**
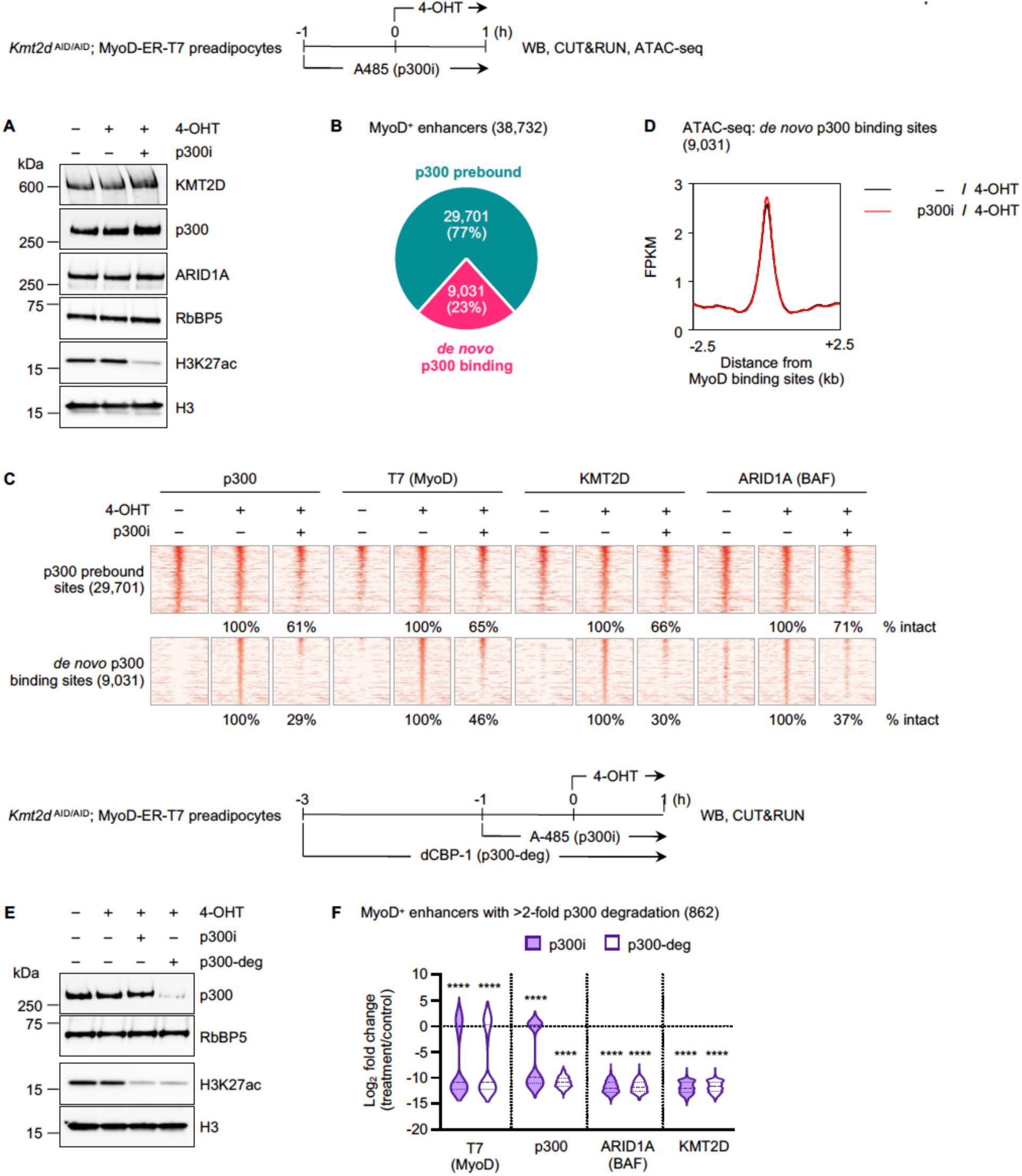
*De novo* binding of MyoD and chromatin modifiers on enhancers depends on p300 acetyltransferase activity. **(A-D)** *Kmt2d* ^AID/AID^; MyoD-ER-T7 preadipocytes were pretreated with p300/CBP inhibitor A-485 (p300i) for 1h, and then 4-OHT was added for 1h to induce MyoD nuclear translocation. Cells were collected for WB, CUT&RUN, and ATAC-seq. **(A)** WB of nuclear extracts for p300, KMT2D, ARID1A (BAF) and histone extracts for H3K27ac. RbBP5 and H3 serve as loading controls. **(B)** Pie chart illustrating p300 binding status on 38,732 MyoD^+^ enhancers. **(C)** Heat maps for CUT&RUN of p300, T7 (MyoD), ARID1A (BAF), and KMT2D on p300 prebound and *de novo* p300 binding sites. Heat maps spanned ± 3kb around MyoD binding sites, and sites were ranked by the intensity of T7 (MyoD) in the 4OHT-treated control. **(D)** Average profiles of normalized ATAC-seq reads on 9,031 *de novo* p300 binding sites with or without p300i treatment. **(E-F)** *Kmt2d* ^AID/AID^; MyoD-ER-T7 preadipocytes were pretreated with p300/CBP degrader dCBP-1 (p300-deg) for 3h or p300i for 1h. Then, 4-OHT was added for 1h to induce MyoD nuclear translocation. **(E)** WB of nuclear extracts for p300 or histone extracts for H3K27ac. **(F)** Violin plot illustrating changes in binding of T7 (MyoD), p300, ARID1A (BAF), and KMT2D upon p300i or p300-deg treatment. The analysis was performed on MyoD^+^ enhancers with >2-fold reduced p300 binding upon p300-deg. Statistical significance was determined using a one-sided Wilcoxon signed-rank test. ****p < 0.0001.

We performed CUT&RUN to examine MyoD, BAF, KMT2D, and p300 binding on enhancers upon p300i treatment. Among the 38,732 MyoD^+^ enhancers defined in Fig. 1D, 77% (29,701 sites) were prebound by p300, and 23% (9,031 sites) showed *de novo* p300 binding (Fig. 4B). p300i treatment reduced MyoD, p300, KMT2D, and BAF binding on approximately 30-40% of p300 prebound sites. In contrast, *de novo* p300 binding sites were more sensitive to p300 inhibition: MyoD enrichment was decreased on 54% of these sites, whereas p300, KMT2D, and BAF binding was reduced on ∼70% of *de novo* p300 binding sites (Fig. 4C). The loss of MyoD and chromatin modifier enrichment was not due to changes in chromatin accessibility upon p300 inhibition (Fig. 4D). These data suggest that p300 might play a more dominant role than previously reported in modulating the binding of LDTFs and chromatin modifiers on enhancers.

We further validated our observations using a heterobifunctional degrader, dCBP-1 (p300-deg), to acutely deplete p300 protein [27]. *Kmt2d* ^AID/AID^; MyoD-ER-T7 cells were pretreated with either p300-deg (3h) or p300i (1h), before MyoD nuclear translocation was induced by 4-OHT (1h). While p300-deg treatment significantly reduced p300 protein levels, p300i treatment did not. However, global H3K27ac levels were decreased by both treatments (Fig. 4E). CUT&RUN analysis revealed that p300-deg and p300i treatment reduced MyoD, BAF, and KMT2D binding on MyoD^+^ enhancers to a similar extent (Fig. 4F). Together, these results indicate that p300, through its acetyltransferase activity, is essential for *de novo* binding of MyoD, BAF, KMT2D, and p300 on enhancers during early stages of myogenic reprogramming.

We also examined the effects of p300 inhibition on endogenous MyoD binding in C2C12 myoblasts, where *de novo* MyoD binding is induced upon differentiation [17]. C2C12 myoblasts were subjected to 2h or 24h of myogenesis, with p300i applied for 2h prior to analysis (Fig. 4-S1A). As expected, a 2h p300i treatment markedly reduced global H3K27ac levels (Fig. 4-S1B). After 2h and 24h of differentiation, we identified 718 and 3,075 *de novo* MyoD^+^ p300^+^ sites, respectively. MyoD motifs were ranked among the most significantly enriched ones (Fig. 4-S1C; S1E). At both time points, inhibition of p300 acetyltransferase activity significantly reduced *de novo* binding of MyoD and p300 on *de novo* MyoD^+^ p300^+^ sites (Fig. 4-S1D; S1F). These findings are consistent with our observations in preadipocytes and demonstrate that p300 enzymatic activity is essential for MyoD and p300 binding on enhancers.

### Cooperative binding of KMT2D, BAF, and p300 on MyoD-intact enhancers

Our results from individual acute perturbations suggest that KMT2D, BAF, and p300 are important for *de novo* MyoD binding on enhancers. To directly compare the effects of KMT2D depletion, BRG1 or p300 inhibition, we focused on a common set of 3,620 MyoD^+^ enhancers that showed *de novo* binding of all three chromatin modifiers and exhibited >2-fold reduction of KMT2D signal upon its depletion (Fig. 5A). All three interventions markedly reduced *de novo* MyoD binding on enhancers, with BRG1 inhibition having the most pronounced impact, followed by p300 inhibition and KMT2D depletion. Acute depletion or inhibition of one chromatin modifier markedly reduced the binding of the other two on MyoD^+^ enhancers (Fig. 5B-C).

**Fig 5.**
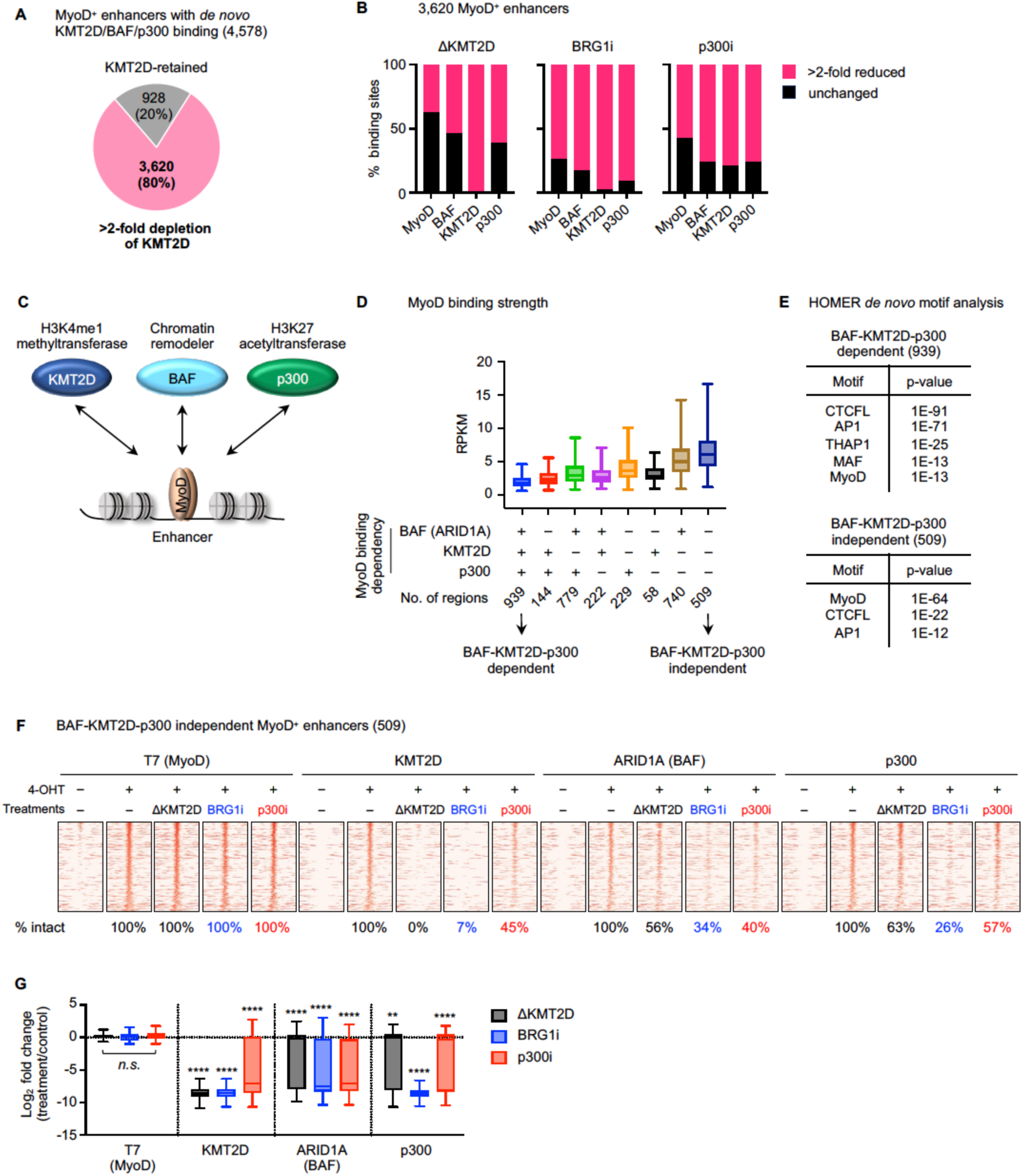
Cooperative binding of chromatin modifiers on MyoD-intact enhancers. **(A)** Pie chart illustrating KMT2D protein depletion on 4,578 MyoD^+^ enhancers with *de novo* binding of KMT2D, BAF, and p300 defined Fig 1F. **(B)** Bar graphs illustrating effects of ΔKMT2D, BRG1i, and p300i on MyoD^+^ enhancers with *de novo* binding of KMT2D, BAF, and p300. **(C)** A model depicting the interdependent relationship between myogenic TF MyoD and chromatin modifiers KMT2D, BAF, p300 on enhancers. **(D)** Box plots showing the normalized MyoD read counts on 3620 MyoD-bound enhancers, grouped by the presence of MyoD binding decrease upon chromatin modifier interventions (MyoD binding dependency). **(E)** HOMER *de* novo motif analysis on BAF-KMT2D-p300 dependent and independent sites defined in (A). **(F-G)** Heat maps **(F)** and corresponding box plots **(G)** illustrating changes in binding of T7 (MyoD), KMT2D, ARID1A (BAF), and p300 on 509 BAF-KMT2D-p300 independent MyoD^+^ enhancers defined in Heat maps spanned ± 3kb around MyoD binding sites, and sites were ranked by the intensity of T7 (MyoD) in the 4OHT control. Statistical significance was determined using a one-sided Wilcoxon signed-rank test. *n.s.* not significant; **p < 0.01; ****p < 0.0001.

Based on the motif analysis of these 3,620 MyoD^+^ enhancers (Fig. 1H and Fig. 5A), MyoD is likely the driving force of chromatin modifier recruitment. Within this set, we further identified two subgroups of sites. One shows >2-fold reduced MyoD binding upon all three perturbations (939 BAF-KMT2D-p300 dependent sites). The other shows intact MyoD binding under all conditions (509 BAF-KMT2D-p300 independent sites) (Fig. 5D). MyoD binding was significantly stronger on independent sites. HOMER motif analysis showed that canonical MyoD motifs were most significantly enriched on independent sites. In contrast, dependent sites exhibited strong enrichment of CTCFL and AP1 motifs, while MyoD motifs ranked lower, suggesting indirect binding (Fig. 5E).

To distinguish direct effects on chromatin modifier enrichment from those secondary to reduced MyoD binding, we focused on 509 BAF-KMT2D-p300 independent MyoD^+^ enhancers. On these 509 enhancers, KMT2D binding was diminished on 93% and 55% of them by BRG1i and p300i treatments, respectively. BAF binding was reduced by KMT2D depletion, BRG1 or p300 inhibition on 44%, 66%, and 60% of sites, respectively. Similarly, p300 binding showed a >2-fold decrease on 37%, 74%, and 43% of sites, upon the loss of KMT2D protein, BRG1 or p300 catalytic activity, respectively (Fig. 5F-G). Interdependent binding of chromatin modifiers was also observed in a broader set of MyoD^+^ enhancers where MyoD binding remained unaffected by individual treatments (Fig. 5-S1).

### GR displays interdependencies with chromatin modifiers on enhancers

To determine whether the dynamics observed between MyoD and chromatin modifiers extend to other TFs, we examined the relationships among the SDTF GR and chromatin modifiers KMT2D, BAF, and p300 following the treatment of synthetic glucocorticoid Dexamethasone (DEX). *Kmt2d* ^AID/AID^ cells were pretreated with 5Ph-IAA for 2h, or with Brg1i or p300i for 1h, and subsequently treated with DEX for 1h to induce GR nuclear translocation. As expected, KMT2D depletion or p300 inhibition markedly reduced KMT2D or H3K27ac levels, respectively, whereas protein levels of p300, KMT2D, and ARID1A, as well as GR nuclear translocation, remained unchanged in all conditions (Fig. 6A).

**Fig 6.**
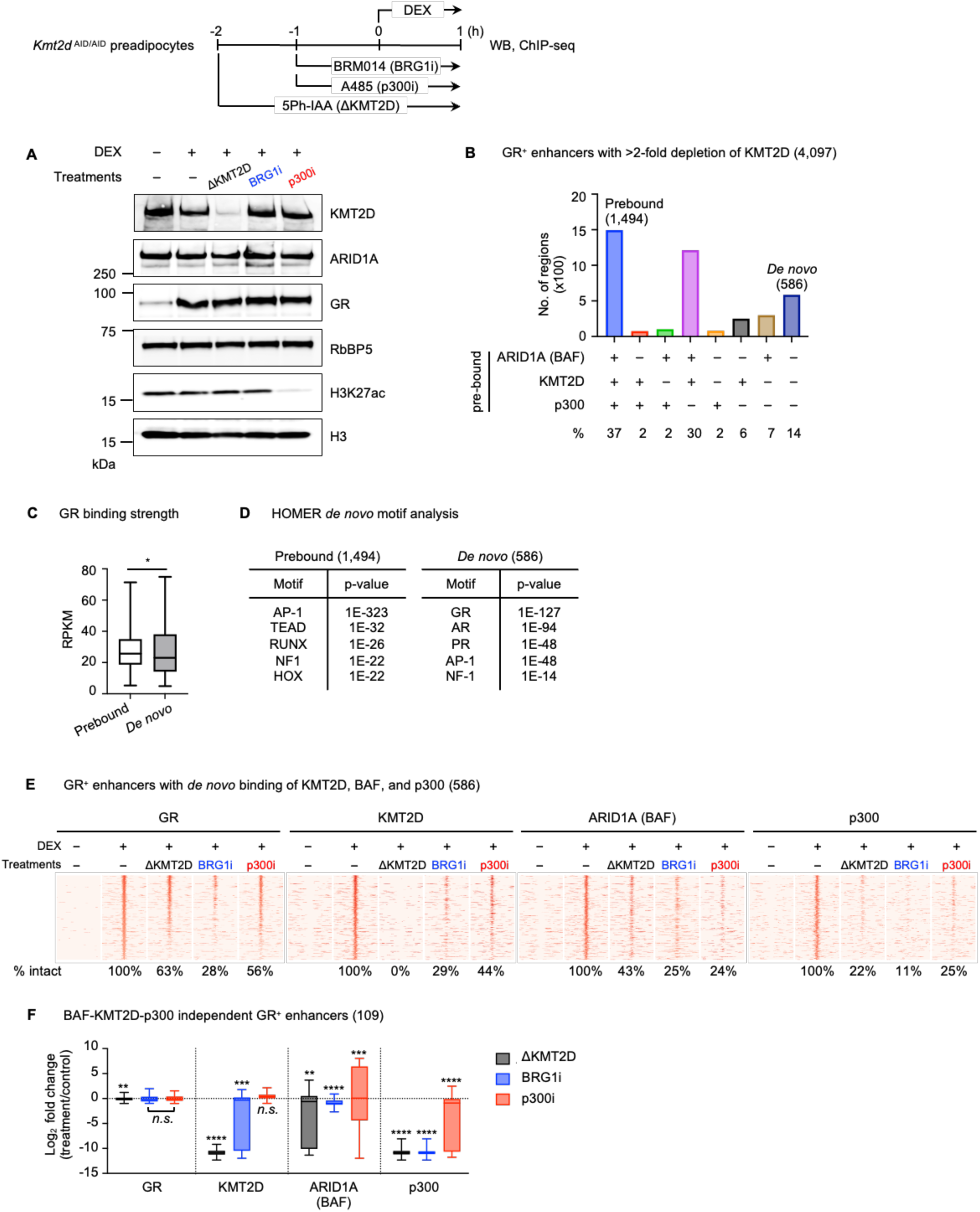
DEX-activated GR displays interdependencies with chromatin modifiers KMT2D, BAF, and p300 on enhancers. *Kmt2d* ^AID/AID^ preadipocytes were pretreated with 5Ph-IAA (ΔKMT2D) for 2h, BRG1i or p300i for 1h, and then 100nM DEX was added for 1h to induce GR nuclear translocation. Cells were collected for WB and ChIP-seq. **(A)** WB of nuclear extracts for KMT2D, ARID1A, and GR and histone extracts for H3K27ac. RbBP5 and H3 serve as loading controls. **(B)** Bar chart showing ARID1A (BAF), KMT2D, and p300 binding on 4,097 GR^+^ enhancers with >2-fold KMT2D depletion, prior to DEX treatment. **(C-D)** Box plots showing the normalized GR read counts **(C)** and HOMER *de* novo motif analysis **(D)** on GR^+^ enhancers with BAF-KMT2D-p300 prebound or *de novo* sites defined in (B). **(E)** Heat maps for ChIP-seq of GR, KMT2D, ARID1A (BAF), and p300 on GR^+^ enhancers with *de novo* binding of BAF, KMT2D, and p300. Heat maps spanned ± 3kb around GR binding sites, and sites were ranked by the intensity of GR in the DEX-treated control. **(F)** Box plots illustrating changes in binding of GR, KMT2D, ARID1A (BAF), and p300 on 109 BAF-KMT2D-p300-independent GR^+^ enhancers. Statistical significance was determined using a one-sided Wilcoxon signed-rank test. *n.s.* not significant; *p < 0.05; **p < 0.01; ***p < 0.001; ****p < 0.0001.

We performed ChIP-seq to examine GR, BAF, KMT2D, and p300 binding following DEX treatment and acute interventions. We identified 4,097 GR^+^ enhancers for analysis, based on the following criteria: (1) GR binding is >2-fold induced upon DEX treatment; (2) BAF, KMT2D, and p300 are enriched; and (3) KMT2D signal is >2-fold decreased upon depletion. Among 4,097 GR^+^ enhancers, 1,494 sites (37%) were prebound by BAF, KMT2D, and p300, while 586 sites (14%) were *de novo* for all three modifiers (Fig. 6B). Unlike our observations with MyoD (Fig. 1G), GR binding strength was comparable between the prebound and *de novo* groups (Fig. 6C). However, GR motifs were only enriched on *de novo* sites, but not on prebound ones (Fig. 6D). We therefore focused our analysis on the 586 *de novo* sites to evaluate the interplay between GR and chromatin modifiers.

Among 586 GR^+^ enhancers with *de novo* binding of KMT2D, BAF, and p300, acute loss of KMT2D protein or BAF and p300 enzymatic activities reduced GR occupancy on 219 (37%), 420 (72%), or 260 (44%) of these sites, respectively (Fig. 6E). These results demonstrate the critical roles of KMT2D, BAF, and p300 in *de novo* binding of GR on enhancers. On these 586 GR^+^ enhancers, KMT2D depletion reduced BAF and p300 binding on 334 (57%) and 459 (78%), respectively. Loss of BAF enzymatic activity also decreased KMT2D, BAF, and p300 binding on 415 (71%), 448 (76%), and 521 (89%) of these sites, respectively. Meanwhile, inhibition of p300 catalytic activity diminished the binding of KMT2D, BAF, and p300 on 328 (56%), 448 (76%), and 440 (75%) of these enhancers, respectively (Fig. 6E).

Among 586 GR^+^ enhancers with *de novo* binding of KMT2D, BAF, and p300, we identified 109 sites that showed less than a 2-fold reduction in GR binding across all three perturbations. We refer to these regions as BAF-KMT2D-p300-independent GR^+^ enhancers (Fig. 6F). On these sites, KMT2D depletion mildly reduced BAF binding but markedly diminished p300 enrichment. Inhibition of BAF enzymatic activity disrupted the binding of all three factors – KMT2D, BAF, and p300 – with p300 being the most impacted. In contrast, p300 inhibition did not affect KMT2D binding and, although significant, caused only modest decreases in BAF and p300 occupancy. Together, these data highlight the interdependent relationships of chromatin modifiers on GR^+^ enhancers (Fig. 6F).

In conclusion, using the LDTF MyoD and the SDTF GR as models and employing acute perturbation of chromatin modifiers, we demonstrate that, in addition to the previously suggested model of sequential recruitment of TFs, KMT2D, BAF, and p300 (Fig. 7A), TFs and these chromatin modifiers also exhibit interdependent binding relationships on enhancers (Fig. 7B).

**Fig 7.**
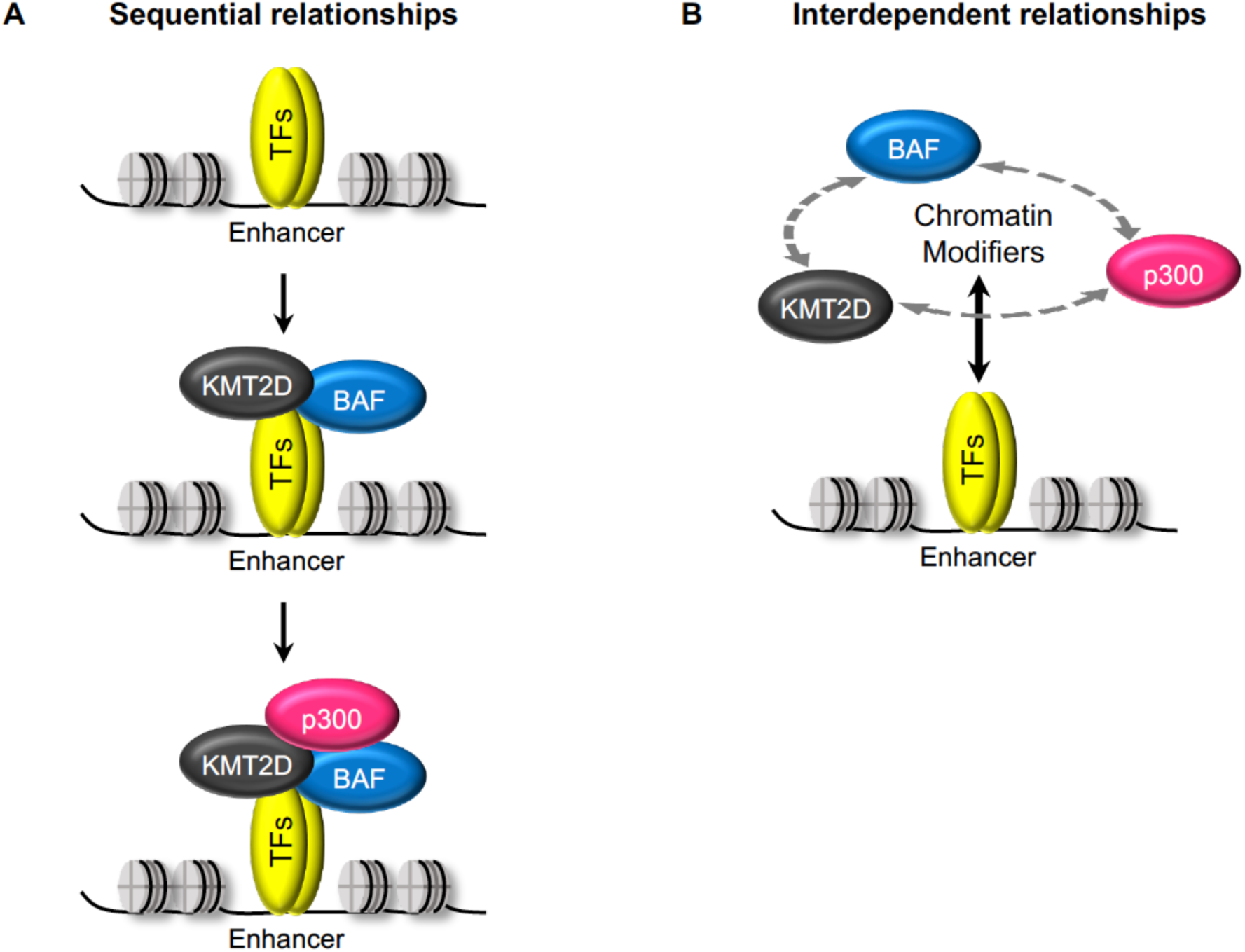
Acute interventions reveal interdependencies among TFs and chromatin modifiers KMT2D, BAF and p300 on enhancers. **(A)** The reported sequential relationships of TF and chromatin modifier enrichment on enhancers established with stable knockout and continuous TF expression models. **(B)** The interdependent relationships between TF and chromatin modifiers KMT2D, BAF, p300 and among chromatin modifiers on enhancers established with acute interventions of chromatin modifiers and inducible nuclear translocation of TFs.

## Discussion

In this study, using robust nuclear translocation systems of MyoD and GR combined with acute depletion of KMT2D protein or short-term inhibition of BAF or p300 enzymatic activity, we illustrated that TFs and chromatin modifiers exhibit interdependent relationships on enhancers. Acute depletion or inhibition of these chromatin modifiers disrupts *de novo* binding of MyoD and GR as well as that of KMT2D, BAF, and p300 on enhancers, with loss of BAF activity having the most pronounced impact. Inhibiting acetyltransferase activity of p300 – a chromatin factor responsible for activating enhancers and thought to function downstream of TFs, KMT2D, and BAF – reveals its critical role in modulating *de novo* TF binding on enhancers. Our data suggest that p300 might have a more prominent role than previously appreciated. Our findings highlight the importance of chromatin modifiers in *de novo* enhancer binding of TFs and indicate the cooperative binding of KMT2D, BAF, and p300 on enhancers.

### Refining KMT2D’s role in TF and chromatin modifier binding with acute protein depletion

Degron-based protein depletion systems offer a powerful approach to study protein functions in living cells and animals. The key advantage lies in their ability to reveal the primary protein functions before the interference of secondary effects or cellular adaptation. Previously, we found that KMT2D knockout reduces C/EBPβ binding on approximately 26% of enhancers [2, 8]. On C/EBPβ-activated enhancers, KMT2D knockout impairs BAF binding, regardless of whether BAF binding is *de novo* or pre-marked [8]. In the current study, results from acute KMT2D depletion experiments not only validated prior observations on TF binding but also distinguished KMT2D’s role on pre-existing and newly established chromatin factor binding: prebound BAF remained stable, while *de novo* BAF binding to MyoD^+^ enhancers was significantly reduced upon the loss of KMT2D (Fig. 2F). This refines and specifies KMT2D’s role in facilitating *de novo* binding of chromatin modifiers alongside with modulating *de novo* binding of TFs.

### BAF activity in modulating *de novo* binding of TF and chromatin modifiers on enhancers

BRG1, through regulating chromatin accessibility, is important for maintaining binding of TFs OCT4 and REST in undifferentiated ESCs [24, 28]. We show that BRG1 activity is required for *de novo* binding of MyoD during myogenic reprogramming of preadipocytes, as well as for DEX-induced GR binding (Fig. 3 & 6). Reduced MyoD binding upon BRG1 inhibition extends beyond a simple decrease in chromatin opening. On MyoD^+^ enhancers with *de novo* BAF binding, loss of BRG1 activity disrupts *de novo* MyoD binding on ∼65% of the sites, while chromatin accessibility is decreased on only 34% of these sites (Fig. 3C & 3E). BRG1 inhibition may alter nucleosome positioning without affecting overall chromatin accessibility, supported by a study reporting that BRG1 depletion disrupts nucleosome organization around key TF motifs in T cells [14]. We observed that BRG1 inhibition reduced ARID1A binding, even when MyoD binding persisted (Fig. 3C & 5F). It is plausible that BRG1 activity is required for maintaining the integrity and/or stability of the BAF complex itself, which in turn reinforces and stabilizes TF binding on these *de novo* enhancers.

Our observations of reduced ARID1A binding upon BRG1 inhibition aligns with CUT&RUN analysis showing a moderate decrease in BRG1 occupancy upon BRG1i treatment in ESCs [29], but contrasts with ChIP-seq data indicating a slight increase [30]. Acute BRG1 protein depletion has been shown to reduce p300 binding in T cells [14], and KMT2D binding on C/EBPβ-bound enhancers [8]. Consistently, upon BRG1 inhibition, both KMT2D and p300 binding is reduced on MyoD^+^ enhancers (Fig. 6C). Together, these findings suggest that BAF enzymatic activity facilitates *de novo* TF binding and the enrichment of BAF complex and other chromatin modifiers on enhancers, partially through promoting chromatin accessibility.

### p300 activity in facilitating *de novo* TF and chromatin modifier binding on enhancers

The potent CBP/p300 inhibitor A-485 enables extensive assessment of the role of CBP/p300 acetyltransferase activity in transcriptional regulation. In ESCs, A-485 treatment results in a significant reduction in global transcription within 30 minutes, without affecting the chromatin binding of pluripotency TFs NANOG or OCT4 [31]. This suggests that CBP/p300 acetyltransferase activity is dispensable for maintaining pre-established TF occupancy. In contrast, we found that p300 inhibition disrupts the *de novo* binding of MyoD on enhancers, an observation also supported by acute p300 degradation in preadipocytes and further confirmed in physiologically relevant C2C12 myoblasts (Fig. 4C & 4F, Fig. 4-S1). Loss of p300 activity also impacts the binding of GR on enhancers (Fig. 7). Our results align more with imaging studies at the *Pou5f1* enhancer, where Sox2 recruits CBP/p300, and CBP/p300 acetyltransferase activity reinforces Sox2 occupancy [32]. In addition, while p300 inhibition does not impact steady-state p300 chromatin binding in ESCs or leukemic MM1.S cells [31, 33], we show that p300 activity is required for *de novo* p300 binding on newly established MyoD^+^ or GR^+^ enhancers.

Despite a reduced enrichment of BAF on enhancers upon p300 inhibition, we found that chromatin accessibility remains intact (Fig. 4D). This is consistent with other reports showing that CBP/p300 inhibitor does not affect chromatin opening [31, 33]. This also indicates that impaired TF binding upon the loss of CBP/p300 enzymatic activity is not due to loss of accessible chromatin. CBP/p300 are known to acetylate many TFs and enhancer-associated chromatin factors [34]. MyoD itself is acetylated on multiple lysine residues [35, 36]. It remains to be determined if impaired TF enhancer binding may result from reduced acetylation of TFs or that of chromatin modifiers that stabilize TF occupancy on enhancers.

In summary, using robust nuclear translocation models of MyoD and GR, combined with acute perturbations of chromatin modifiers, our study demonstrates that KMT2D, BAF, and p300 are active, interdependent facilitators of *de novo* TF binding on enhancers. These chromatin modifiers also modulate each other’s binding on TF-bound enhancers. Future studies using higher temporal resolution techniques, such as live-cell imaging or optogenetics, may help define the precise order, kinetics, and relationships of chromatin modifier recruitment during the initiation of enhancer activation.

## Materials and Methods

### DNA constructs/Plasmids

MyoD-ER(T) cDNA was PCR amplified from pLv-CMV-MyoD-ER(T) plasmid (#26809, Addgene) [20] and subcloned into pCW57.1puro-MCS-3xT7 [37] to generate MyoD-ER(T)-3xT7. MyoD-ER(T)-3xT7 cDNA was further subcloned into pWZLhygro (#18750, Addgene) to generate pWZLhygro-MyoD-ER(T)-3xT7. pWZLhygro-MyoD-K/R-ER(T)-3xT7 where K99, K102, and K104 of MyoD were mutated to R was generated from pWZLhygro-MyoD-ER(T)-3xT7 with Q5^®^ Site-Directed Mutagenesis Kit (#E0554S, NEB). pBabepuro-OsTIR1(F74G)-9xMyc was generated from pBabepuro-OsTIR1-9xMyc (#80074, Addgene) by site-directed mutagenesis. All plasmids were confirmed by Sanger DNA sequencing.

### Antibodies and chemicals

Anti-T7 (D9E1X, #13246), anti-ARID1A/BAF250A (D2A8U, #12354), and anti-p300 (D8Z4E, #86377) were from Cell Signaling Technology. Rabbit control IgG (#13-0042K) and anti-H3K4me1 (#13-0040) were from EpiCypher. Anti-H3K27ac (ab4729) was from Abcam. Anti-p300 (C-20, sc-585), anti-MyoD (5.8A, sc-32758), and anti-GR (M-20, sc-10004) were from Santa Cruz. The homemade anti-KMT2D has been previously described [38].

BRG1 Inhibitor BRM014 (#HY-119374) and 5Ph-IAA (#HY-134653) were from MedChemExpress (MCE) and used at 1µM. PROTAC p300/CBP degrader dCBP-1 (#HY-134582) from MCE was used at 250nM. p300/CBP inhibitor A-485 (#6887) was from Tocris Bioscience and used at 3µM. (Z)-4-Hydroxytamoxifen (4-OHT) (#H7904) and Dexamethasone (DEX) (#D4902) were from Millipore-Sigma and used at 400nM and 100nM, respectively.

### Mouse strains and housing

To generate *Kmt2d* ^AID/AID^ mice, a 132bp DNA fragment encoding AID tag was inserted after the start codon located on exon 2 of *Kmt2d* by CRISPR/Cas9-mediated homology-directed repair. Successful integration of AID tag was confirmed with genotyping and Sanger DNA sequencing. *Kmt2d* ^AID/AID^ mice were crossed with *Kmt2c* ^f/f^; *Cre-ER* mice [22] to obtain *Kmt2c* ^f/f^; *Kmt2d* ^AID/AID^; *Cre-ER* mice. Mice were housed in a room with a controlled temperature of 22°C and humidity of 45-65% under a 12h light and 12h dark cycle. All mouse work was approved by the Animal Care and Use Committee of NIDDK, NIH.

### Preadipocyte cultures and myogenesis assay

Primary brown preadipocytes were isolated from interscapular brown adipose tissue of newborn pups and were immortalized by infection with retroviruses carrying pBABE-neo large T cDNA expressing SV40T as described [39]. Immortalized preadipocytes were sequentially infected with retroviruses expressing pBabepuro-OsTir1(F74G)-9xMyc and pWZLhygro-MyoD-ER(T)-3xT7. The immortalized preadipocytes were routinely cultured in growth media: DMEM (#01-013-CV, Corning) supplemented with 10% Foundation FBS (#SH30910, Cytiva). C2C12 myoblasts were cultured in DMEM supplemented with 20% Foundation FBS.

For myogenesis assay, preadipocytes and C2C12 were grown in growth media for 2 days. On the day of experiments, growth medium was replaced with differentiation medium: DMEM supplemented with 2% horse serum (#H1270, Millipore-Sigma). For differentiation assay that lasts more than 48h, the media was changed after 48h and every two days thereafter.

To induce MyoD-ER-T7 and GR translocation, 4-OHT and DEX was used at the final concentration of 400nM and 100nM, respectively. BRM014 and 5Ph-IAA were used at the final concentration of 1µM to inhibit BRG1 chromatin remodeling activity and deplete KMT2D, respectively, while A-485 was used at 3µM to inhibit CBP/p300 acetyltransferase activity.

### Western blot

Nuclear proteins were extracted from preadipocytes with a high-salt method. Briefly, cells were washed with cold PBS, resuspended in buffer A (10mM HEPES, pH 7.9, 1.5mM MgCl_2_, 10mM KCl, and 0.1% NP-40) supplemented with protease inhibitors (PI’s, #04693159001, Roche), and incubated on ice for 10 minutes (mins). Nuclei were harvested by 10-min centrifugation at 2,000rpm. Nuclear proteins were extracted in buffer C (20mM HEPES pH 7.9, 1.5mM MgCl_2_, 420mM NaCl, 0.2mM EDTA, and 25% glycerol, supplemented with PI’s) for 20 mins on ice. Nuclear extracts were separated using 4-15% Tris-Glycine Gradient Gels (#4561085, Bio-Rad) or 3-8% NuPAGE™ Tris-Acetate Mini Protein Gel (#EA0375BOX, ThermoFisher). Proteins were transferred to PVDF membranes, using the iBlot 2 Gel Transfer Device (Invitrogen) or wet transfer method. The membranes were probed with specific antibodies.

Histones were extracted with the following protocol. Nuclei were first isolated from preadipocytes with Triton Extraction Buffer (PBS containing 0.5% Triton X-100 and PI’s), on ice for 10 mins. Histones were extracted with 0.2N HCl on ice for >2h. The histone-containing supernatants were then neutralized with 10% 2M NaOH (v/v). Histones were separated using 4-15% Tris-Glycine Gradient Gels and transferred to PVDF membranes using the Trans-Blot Turbo Transfer System (Bio-Rad). The membranes were probed with specific antibodies.

### RNA isolation and qRT-PCR analysis

Total RNA was extracted using TRIzol (#15596018, Invitrogen) and reverse transcribed using ProtoScript II first-strand cDNA synthesis kit (#E6560, NEB), following manufacturer’s instructions. Quantitative RT-PCR (qRT-PCR) was performed in triplicates with the Luna® Universal qPCR Master Mix (#M3003, NEB) using QuantStudio 5 Real-Time PCR System (Applied Biosystems™). PCR amplification parameters were 95°C (3 mins) and 40 cycles of 95°C (15 sec), 60°C (60 sec), and 72°C (30 sec).

Statistical significance was calculated using the two-tailed unpaired *t*-test on two experimental conditions. qRT-PCR of *Myog* and *Ckm* was done using the following primers: *Myog*: forward 5’-CCCTGCCCCACAGGGGCTGTG-3’ and reverse 5’-ACGCCACAGAAACCTGAGCCC-3’; *Ckm*: forward 5’-CGCCAGCTAGACTCAGCACT-3’ and reverse 5’-CCCTGCGAGCAGATGAGCTT-3’

### CUT&RUN

CUT&RUN was performed using CUTANA™ ChIC/CUT&RUN Kit (#14-1048, Epicypher, version 4), following the manufacturer’s instructions with modifications. 0.5×10^6^ preadipocytes were used for each CUT&RUN reaction. Cells were gently fixed with 0.1% formaldehyde for 1 min at room temperature (RT). After two PBS washes, cells were scraped out in PBS and harvested via centrifugation at 600g for 3 mins. Cells were resuspended in 100µl cold Nuclei Extract Buffer (20mM HEPES pH 7.9, 10mM KCl, 0.1% Triton X-100, 20% Glycerol, 1mM MgCl_2_ supplemented with PI’s and 0.5mM Spermidine) and lysed on ice for 10 mins. Nuclei were then harvested via centrifugation, resuspended in cold Nuclei Extract Buffer, and incubated with activated 10µL Concanavalin A beads for 10 mins at RT. The cell-beads were captured by a magnetic stand and were incubated with the desired primary antibody in 50µL of cold Antibody buffer (wash buffer plus 0.01% Digitonin and 2mM EDTA) overnight at 4°C. Any unbound antibody was washed twice with 200µL cold Cell Permeabilization buffer (wash buffer plus 0.01% Digitonin). The supernatant was discarded via bead liquid separation using a magnetic stand. 2.5µL pAG-MNase was added, and the complex was incubated for 10 mins at RT. The bound complexes were then washed twice with 200µL cold Cell Permeabilization buffer. To activate pAG-MNase, 100mM CaCl_2_ was added to the bound complexes. The digestion reaction was carried out at 4°C for 2h and neutralized by the addition of STOP Buffer Master Mix including 1ng *E. coli* Spike-in DNA, followed by an incubation at 37°C for 10 mins to release cleaved DNA fragments. The released DNA was separated out using a magnetic stand. The supernatant containing CUT&RUN enriched DNA was purified via a spin column.

CUT&RUN enriched DNA was used for library construction. Library construction was performed using the NEBNext Ultra™ II DNA Library Prep Kit for Illumina (#E7645L, NEB) and NEBNext Multiplex Oligos for Illumina (#E6440L, NEB). All reaction volumes were reduced to half of the recommended reaction volume. Adapter ligation was performed using the NEB Adapter stock diluted by 10-fold. Post adapter ligation, 1µl NEB USER enzyme was added to the reaction and incubated at 37°C for 15 mins. After adapter ligation and hairpin cleavage, 1X AMPure XP Beads (#A63881, Beckman Coulter) were added to the reaction to recover DNA fragments. Purified DNA was amplified with the NEBNext Multiplex Oligos. To avoid amplification of longer DNA fragments, the following PCR cycling parameters were used: 98°C (45 sec); 14 cycles at 98°C (15 sec) and 60°C (10 sec); and 98°C (60 sec). The amplified libraries were purified with 1X AMPure XP Beads. The libraries were paired-end sequenced on the Illumina NovaSeqX or NextSeq2000 System.

### ChIP-seq

Chromatin immunoprecipitation (ChIP) was performed as previously described [8]. In summary, cells were cross-linked with 1-2% formaldehyde for 10 minutes and then quenched using 125 mM glycine at RT. The cells were washed with cold PBS twice, and nuclei were harvested in Farnham lysis buffer (5mM PIPES (pH 7.5), 85mM KCl, 1% NP-40, supplemented with PI’s). Nuclei were then resuspended in TE (10mM Tris-Cl pH7.6 and 1mM EDTA) and sonicated for 17 min (30 s on/off cycle). Lysates were supplemented with detergents to make 1X RIPA buffer (10mM Tris-Cl pH 7.6, 1mM EDTA, 0.1% SDS, 0.1% sodium deoxycholate, 1% Triton X-100, PI’s) and centrifuged at 13,000 g for 10 minutes at 4°C to remove debris. The supernatant containing sheared DNA was transferred to a new tube for ChIP. Each ChIP reaction included 4 to 10µg of target antibodies, 20ng of spike-in chromatin (#53083, Active Motif), and 2µg of spike-in antibody (anti-H2Av, #61686, Active Motif), incubated on a rotator at 4°C overnight. ChIP samples were then mixed with 50 μL of prewashed Protein A Dynabeads (#10002D, ThermoFisher) and incubated for 3 hours at 4°C. The beads were collected using a magnetic rack and washed twice with RIPA buffer, RIPA buffer containing 300 mM NaCl, cold LiCl buffer, and PBS. ChIP and input samples were added to 100μL of ChIP elution buffer containing 0.1M NaHCO3, 1% SDS, and 20μg of Proteinase K, and incubated at 65°C overnight. ChIP DNA was purified using the QIAquick PCR purification kit (#28104, Qiagen). Entire ChIP DNA or 300 ng of the input DNA were used to construct libraries with NEBNext Ultra™ II DNA Library Prep Kit for Illumina, NEBNext Multiplex Oligos for Illumina, and AMPure XP Beads. The libraries were paired-end sequenced on the Illumina NovaSeqX System.

### ATAC-Seq

The Assay for Transposase-Accessible Chromatin with high-throughput sequencing (ATAC-Seq) was performed using ATAC-Seq Kit (#53150, Active Motif), following the manufacturer’s protocols. In short, 100,000 cells were freshly collected for each ATAC reaction. After one PBS wash, cells were resuspended in 100µl ice-cold ATAC lysis buffer and immediately spun down at 500g for 10 mins at 4°C. Cell pellet was then incubated in 50µL Tagmentation Mix at 37°C for 30 mins in a thermomixer set at 800rpm. Tagmented DNA was then purified via a spin column and amplified using i7- and i5-index primers provided in the kit. The amplified library was purified with 1.2X SPRI bead solution, following the kit instruction. Samples were paired-end sequenced on the Illumina NextSeq2000 System.

### Computational analysis

#### CUT&RUN, ChIP-seq, ATAC-seq data processing

Raw sequencing data were aligned to the mouse mm10 genome (CUT&RUN, ChIP-seq, and ATAC-seq), the *E. coli* genome eck12 (CUT&RUN), and *Drosophila* genome dm6 (ChIP-seq) using bowtie2 (v.2.3.4.1).

All datasets were normalized using spike-in read counts prior to downstream analyses. For CUT&RUN, normalization factors were determined in two steps. First, the ratio of *E. coli* eck12 reads to mouse mm10 reads was calculated for each sample. Second, the normalization factors were obtained by dividing the lowest ratio of eck12/mm10 by the ratio of each corresponding sample. For ChIP-seq data, normalization factors were determined by dividing the lowest *Drosophila* dm6 reads by the dm6 reads of each corresponding sample. Normalized read counts for all datasets were adjusted by multiplying the read counts by their respective normalization factors.

#### CUT&RUN and ChIP-seq peak calling

Following alignment, peaks were identified using the SICER2 algorithm [https://zanglab.github.io/SICER2/] [40]. For peak calling of H3K4me1 and H3K27ac, we used the window size of 200bp, the gap size of 200bp, and the false discovery rate (FDR) of 1E-3. For peak calling of non-histone factors, we used the window size of 50bp, the gap size of 50bp, and the FDR of 1E-10. An effective genome fraction of 0.8 was applied to all datasets. All other parameters were set to default values.

#### Motif analysis

To identify TF binding motifs enriched within the genomic regions of interest, we used the HOMER motif discovery algorithm [http://homer.ucsd.edu/homer/] [41], executing the findMotifsGenome.pl command with default parameters. From the knownResults and *de novo* results output, we selected and presented representative TFs from each family that share the same consensus sequence to avoid redundancy.

#### Clustering of MyoD-ER binding regions

To investigate the dynamics of MyoD-ER genomic binding across time points after 4-OHT treatment, read counts were calculated on the union of MyoD-ER sites (Fig. S1). Normalization was applied using two methods: (1) scaling by total reads on peaks and (2) applying spike-in normalization factors derived from the ratio of *E. coli* Eck12 reads to mouse mm10 reads. Z-scores were calculated for each genomic region by subtracting the mean read count across the time points and dividing by the standard deviation. These Z-scores were used for K-means clustering and visualized as heatmaps using the gplots package in R.

#### Heat maps, box plots, violin plots and average profiles

Heat maps were generated using custom Python scripts that processed matrix files with 50 bp resolution and visualized with Seaborn, with customizable color palettes for data representation. To enable comparative analysis across experimental groups, enhancer regions were ordered by signal intensity of MyoD-ER at the center of a 400bp window under control conditions (4OHT) and log-transformed using the log2(x+1) scale. Box plots were used to display the normalized MyoD-ER or GR read counts (Fig. 1, 5, 6). Box plots were also used to illustrate the ratios of normalized ARID1A (BAF) and p300 read counts between KMT2D-depleted and control cells (Fig. 2) or the ratios of normalized T7 (MyoD), KMT2D, ARID1A (BAF), and p300 read counts between KMT2D-deleted, BRG1 or p300 inhibition and control cells (Fig. 5 & 6). Each box plot shows the first and third quartiles, with a horizontal line indicating the median and whiskers representing the minimum and maximum values. Violin plots were used to visualize the ratios of normalized read counts between p300i- or p300-degrader-treated and control cells, also showing the first and third quartiles and the median (Fig. 4). Outliers were not included. Statistical significance was assessed using a one-sided Wilcoxon signed-rank test or a two-sided, unpaired Mann Whitney test, as indicated in each figure legend. Average profiles were generated using normalized read counts binned at 5bp intervals, spanning from the center of MyoD-ER binding sites to 2.5kb upstream and downstream (Fig. 3 & 4).

#### Chromatin accessibility on MyoD^+^ enhancers

Chromatin accessibility on MyoD^+^ enhancers was quantified by calculating ATAC-seq read counts before and after MyoD nuclear translocation (Fig. 3). Three groups of enhancers were defined based on accessibility changes and normalized read counts (RPKM) in MyoD-translocated condition. Constitutively open enhancers were defined as regions showing less than a 2-fold increase in ATAC-seq signal after MyoD translocation and exhibiting accessibility of RPKM > 1. MyoD-dependent opening enhancers were identified as regions showing more than a 2-fold increase in ATAC-seq signal upon MyoD translocation, also with RPKM > 1. Regions with low overall accessibility, defined by RPKM < 1, were designated as low-opening enhancers.

## Data Availability

The datasets supporting this study are available from the corresponding author upon reasonable request. All ChIP-seq, CUT&RUN, and ATAC-seq data have been deposited in the NCBI Gene Expression Omnibus (GEO) under accession number GSE290849.

## Acknowledgements

We thank the NHLBI DNA Sequencing and Genomics Core and the NIAMS Genomic Technology Section for next-generation sequencing.

This work was supported by the Intramural Research Program of NIDDK, NIH to K.G.

## Author Contributions

Conceptualization, H.T.V., J.-E.L., and K.G.; Methodology, H.T.V., J.-E.L., Y.-K.P., C.L., S.I., S.D., V.S, and W.P.; Investigation, H.T.V., J.-E.L.; Software, Formal Analysis, and Data Curation, H.T.V., J.-E.L.; Writing – Original Draft, H.T.V., J.-E.L.; Writing – Review & Editing, H.T.V., J.-E.L., K.G.; Project Administration and Funding Acquisition, K.G.

## Disclaimers

This research was supported by the Intramural Research Program of the National Institute of Diabetes and Digestive and Kidney Diseases (NIDDK) and the National Institute of Arthritis and Musculoskeletal and Skin Diseases (NIAMS) within the National Institutes of Health (NIH). The contributions of the NIH author(s) are considered Works of the United States Government. The findings and conclusions presented in this paper are those of the author(s) and do not necessarily reflect the views of the NIH or the U.S. Department of Health and Human Services.

---

**Fig 1-S1.**
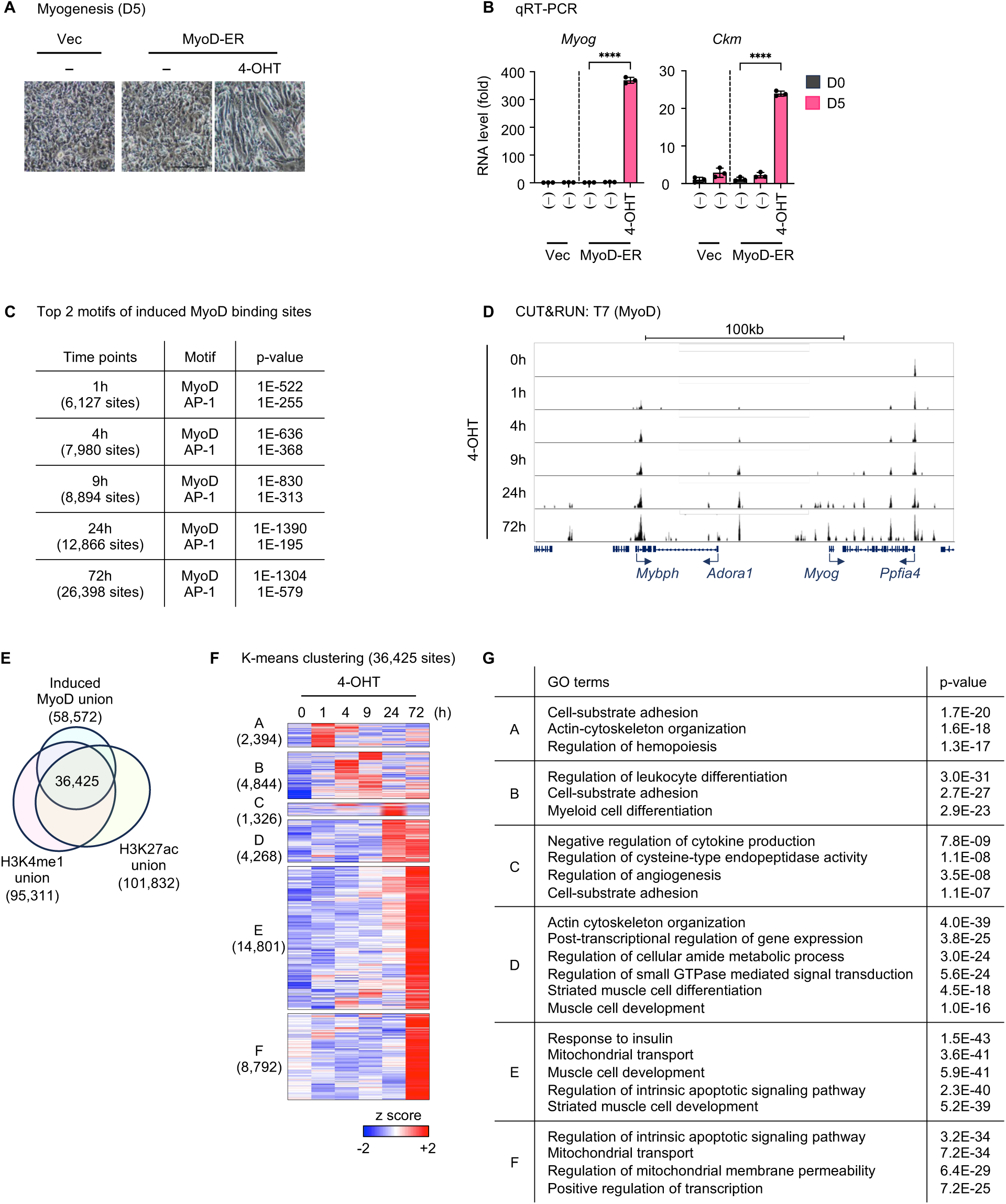
Dynamic binding of nuclear translocated MyoD during myogenesis. **(A)** Cell morphology under microscope at day 5 (D5) of myogenesis. **(B)** qRT-PCR analysis of myogenic markers *Myog* and *Ckm* at D0 and D5 of myogenesis. Data represents mean ± SD, **** p ≤ 0.0001 **(C-G)** MyoD-ER expressing preadipocytes were treated with 4-OHT during myogenesis. Cells were harvested for CUT&RUN at indicated time points. **(C)** HOMER motif analysis of induced MyoD binding sites. All regions were used, and only the top 2 motifs are shown. **(D)** Genome browser view of MyoD binding around the *Myog* locus. **(E)** Venn diagrams depicting 36,425 4-OHT-induced MyoD binding sites used for k-means clustering. **(F)** Heat maps of K-means clustering depicting MyoD binding dynamics on 36,425 sites defined in (E). **(G)** GO analysis of genes associated with MyoD binding sites in each cluster defined in (F).

**Fig 2-S1.**
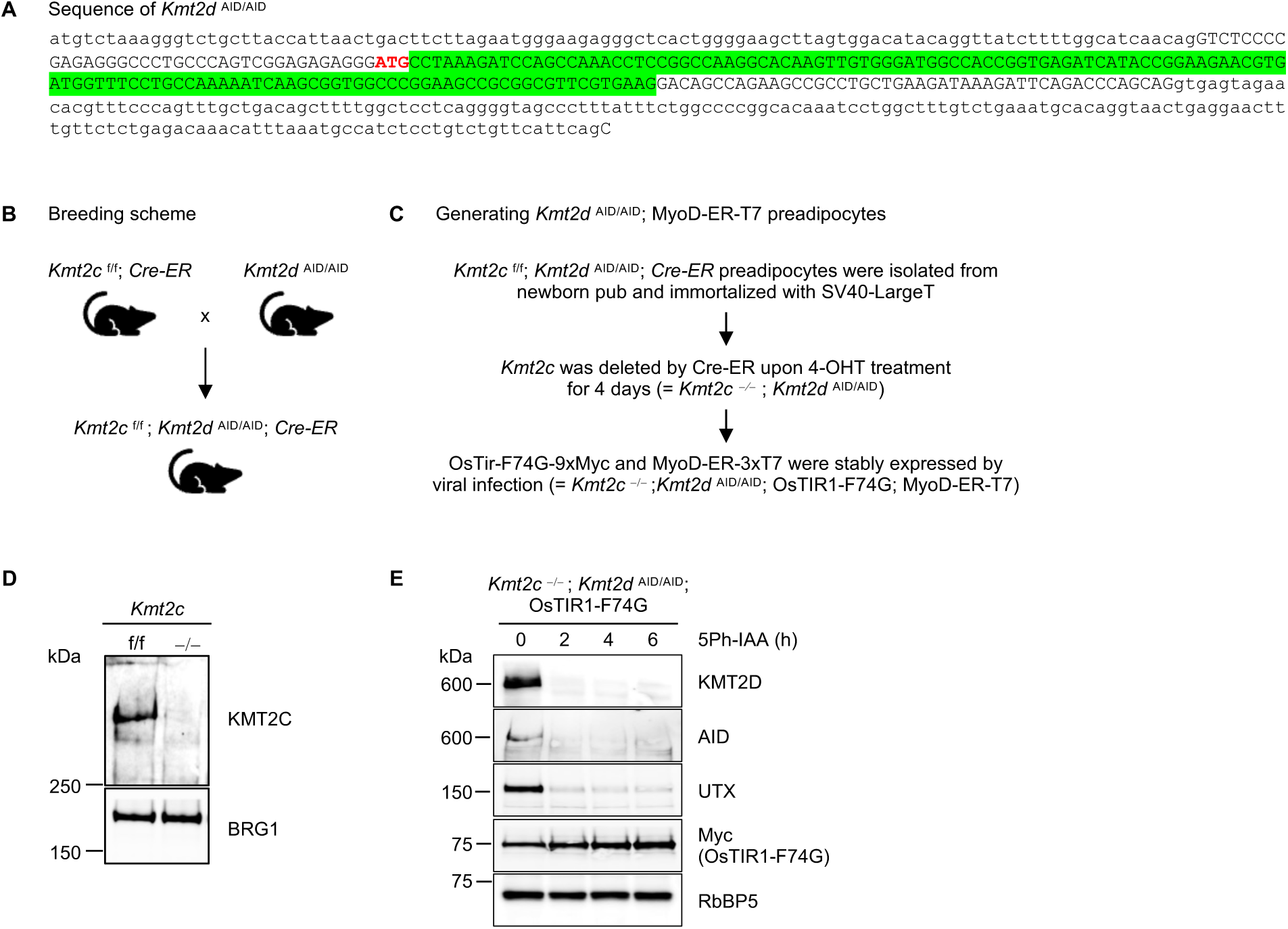
Generation of *Kmt2c* ^-/-^*; Kmt2d* ^AID/AID^ preadipocytes expressing OsTIR1-F74G-Myc and MyoD-ER-T7. **(A)** Sanger sequencing results verifying the integration of 132-bp AID sequence after the start codon (ATG) of *Kmt2d*. **(B)** Breeding scheme for generating *Kmt2c* ^f/f^*; Kmt2d* ^AID/AID^; *Cre-ER* mice. **(C)** Diagram illustrating the workflow to generate *Kmt2c* ^-/-^*; Kmt2d* ^AID/AID^ preadipocytes expressing OsTIR1-F74G-Myc and MyoD-ER-T7. **(D)** WB of nuclear extracts confirming KMT2C knockout. Antibodies used were indicated on the right. BRG1 was a loading control. **(E)** WB of nuclear extracts confirming the depletion of KMT2D in *Kmt2d* ^AID/AID^; MyoD-ER-T7 preadipocytes upon 5Ph-IAA treatment for various time durations. Antibodies used were indicated on the right. RbBP5 was a loading control.

**Fig 4-S1.**
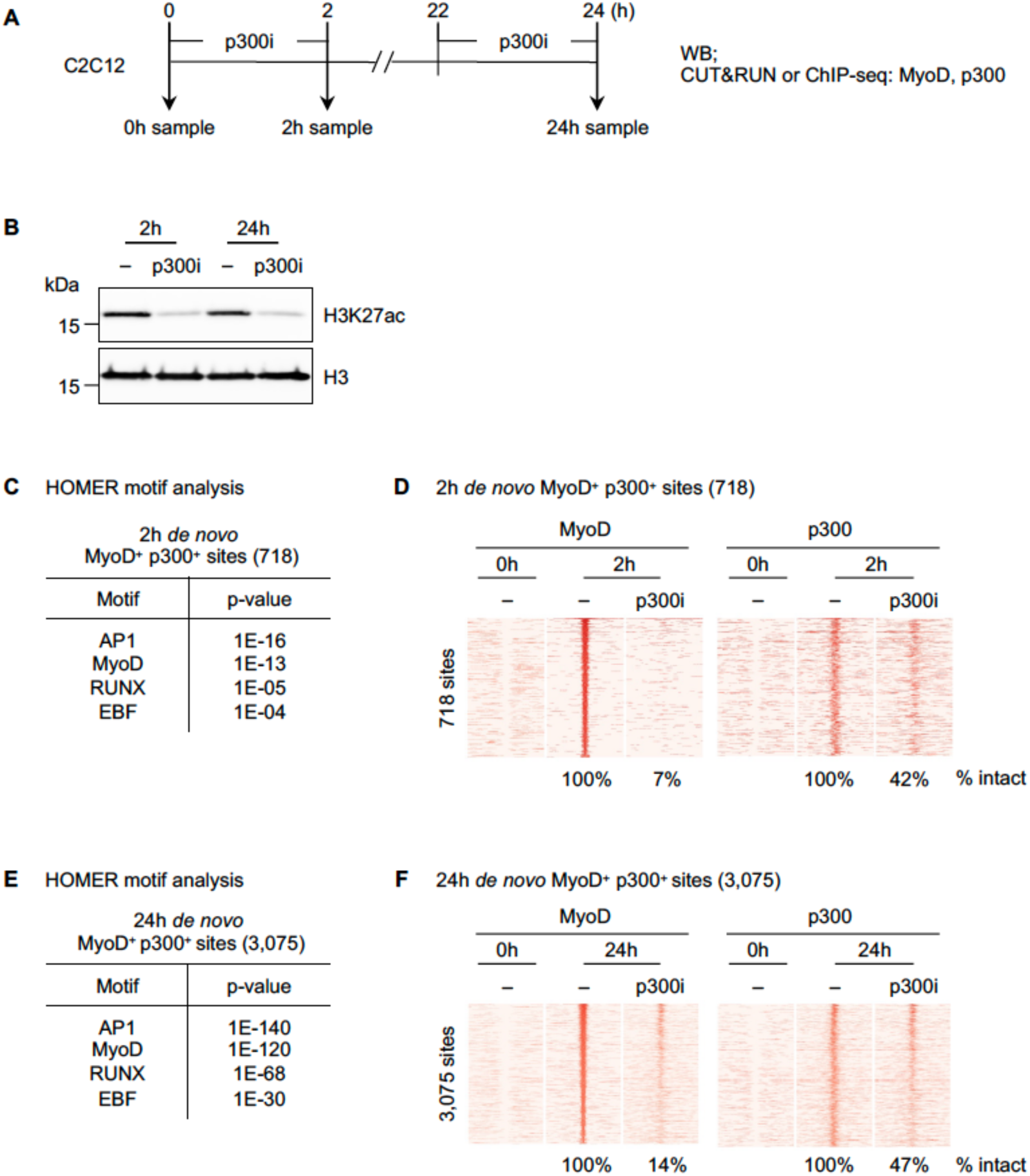
p300 enzymatic activities are required for *de novo* MyoD binding on enhancers in C2C12. **(A)** C2C12 myoblasts were subjected to 2h or 24h of myogenesis and 2h of p300 inhibition, with A-485 (p300i) applied for 2h prior to experiments. **(B)** WB of histone extracts for H3K27ac. H3 serves as the loading control. **(C-D)** Homer motif analysis **(C)** and heat maps for ChIP-seq of MyoD and CUT&RUN of p300 **(D)** on 718 *de novo* MyoD^+^ p300^+^ sites after 2h of differentiation. **(E-F)** Homer motif analysis **(E)** and heat maps for ChIP-seq of MyoD and CUT&RUN of p300 **(F)** on 3,075 *de novo* MyoD^+^ p300^+^ sites after 24h of differentiation.

**Fig 5-S1.**
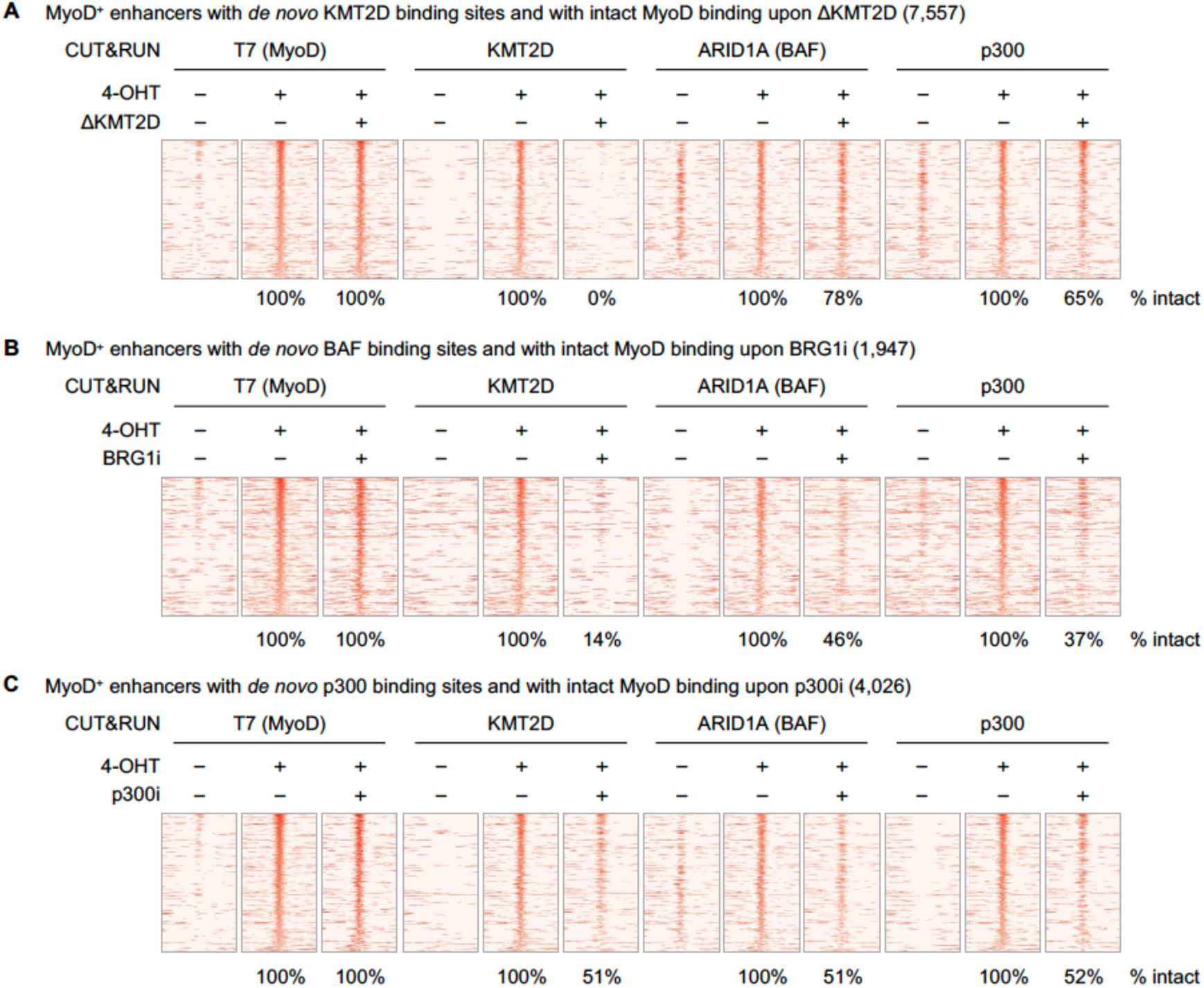
Chromatin modifiers KMT2D, BAF, and p300 regulate each other’s binding on MyoD-intact enhancers. **(A-C)** Heat maps for CUT&RUN data on MyoD^+^ enhancers with *de novo* KMT2D binding and with intact MyoD signals upon ΔKMT2D **(A)**, on MyoD^+^ enhancers with *de novo* BAF binding and with intact MyoD signals upon BRG1i **(B)**, and on MyoD^+^ enhancers with *de novo* p300 binding and with intact MyoD signals upon p300i. All heat maps spanned ± 3kb around MyoD binding sites, and sites were ranked by the intensity of T7 (MyoD) in the 4OHT-treated control.

**Fig 7-S1.**
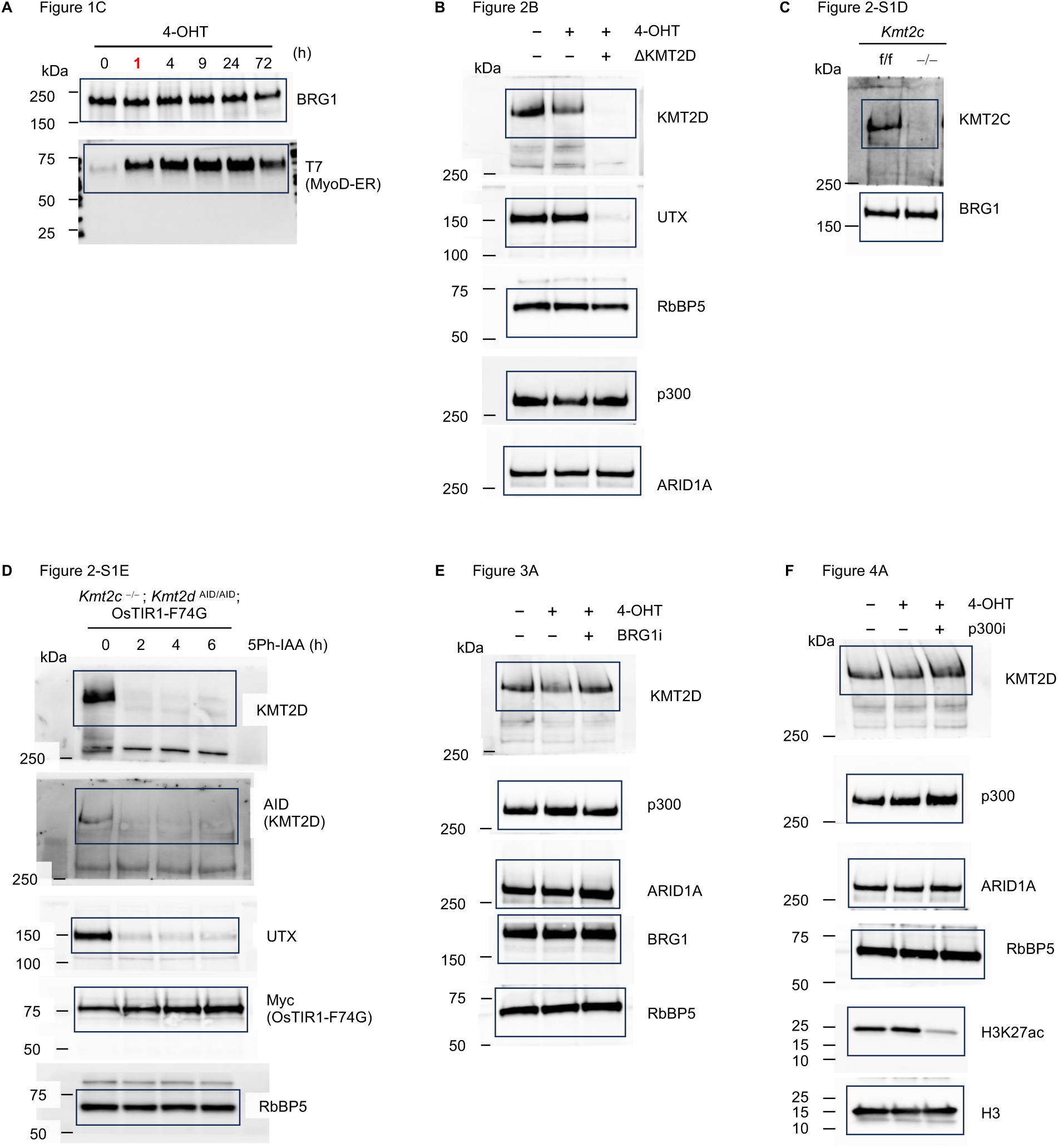
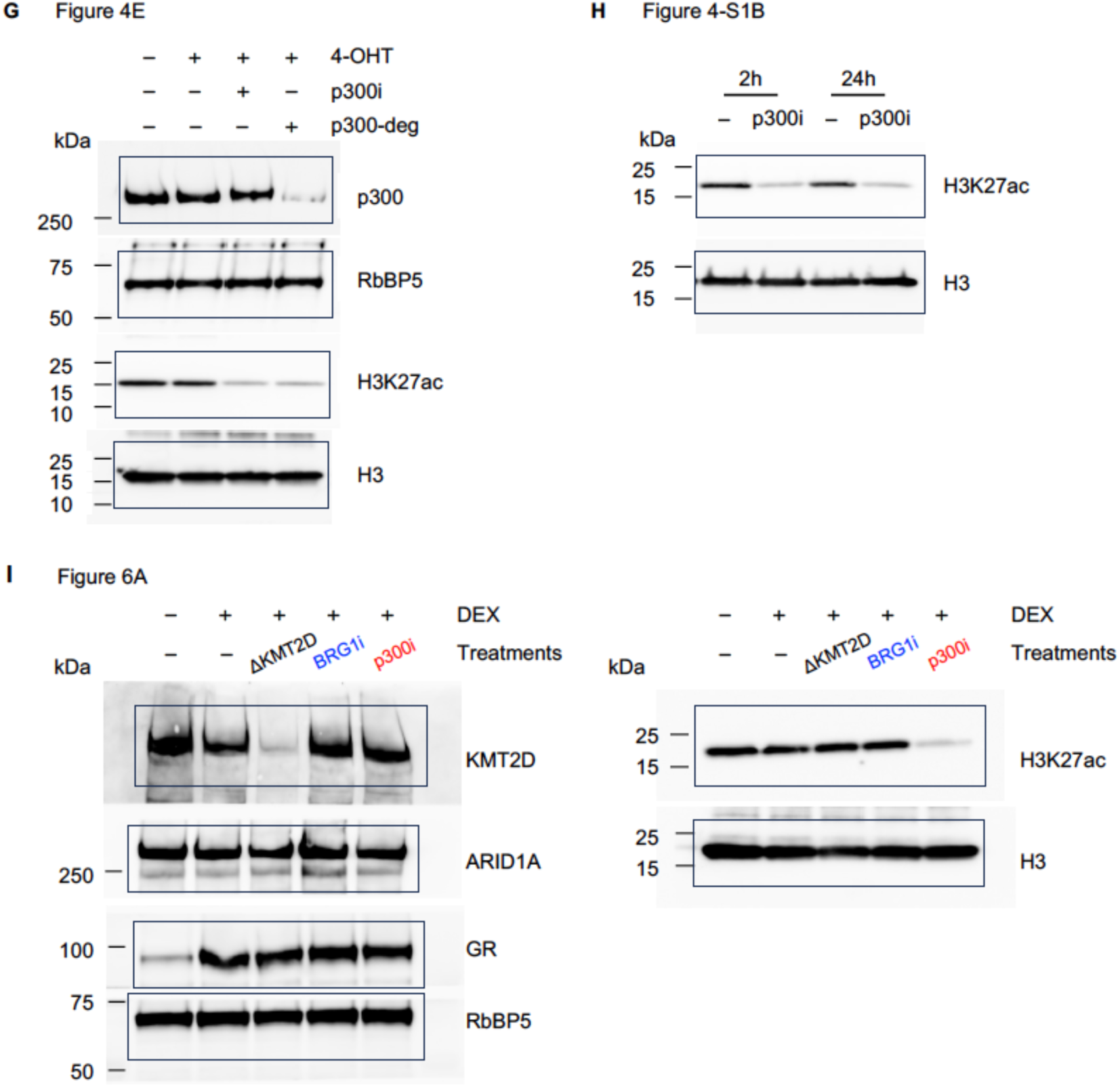
Unprocessed WB images. Uncropped Western blots related to Fig. 1C **(A)**, Fig. 2B **(B)**, Fig. 2-S1D **(C)**, Fig. 2-S1E **(D)**, Fig. 3A **(E)**, Fig. 4A **(F)**, Fig. 4E **(G)**, Fig. 4-S1B **(H)**, and Fig. 6A **(I**).

## References

1. Heinz, S., et al., The selection and function of cell type-specific enhancers. Nat Rev Mol Cell Biol, 2015. 16(3): p. 144–54.

2. Lee, J.E., et al., H3K4 mono- and di-methyltransferase MLL4 is required for enhancer activation during cell differentiation. Elife, 2013. 2: p. e01503.

3. Wang, C., et al., Enhancer priming by H3K4 methyltransferase MLL4 controls cell fate transition. Proc Natl Acad Sci U S A, 2016. 113(42): p. 11871–11876.

4. Jin, Q., et al., Distinct roles of GCN5/PCAF-mediated H3K9ac and CBP/p300-mediated H3K18/27ac in nuclear receptor transactivation. EMBO J, 2011. 30(2): p. 249–62.

5. Visel, A., et al., ChIP-seq accurately predicts tissue-specific activity of enhancers. Nature, 2009. 457(7231): p. 854–8.

6. Xu, L., H. Xuan, and X. Shi, Dysregulation of the p300/CBP histone acetyltransferases in human cancer. Epigenomics, 2025. 17(3): p. 193–208.

7. Clapier, C.R., et al., Mechanisms of action and regulation of ATP-dependent chromatin-remodelling complexes. Nat Rev Mol Cell Biol, 2017. 18(7): p. 407–422.

8. Park, Y.K., et al., Interplay of BAF and MLL4 promotes cell type-specific enhancer activation. Nat Commun, 2021. 12(1): p. 1630.

9. Van, H.T., et al., KMT2 Family of H3K4 Methyltransferases: Enzymatic Activity-dependent and - independent Functions. J Mol Biol, 2024. 436(7): p. 168453.

10. Hota, S.K. and B.G. Bruneau, ATP-dependent chromatin remodeling during mammalian development. Development, 2016. 143(16): p. 2882–97.

11. Mendiratta, G., et al., Cancer gene mutation frequencies for the U.S. population. Nat Commun, 2021. 12(1): p. 5961.

12. Lai, B., et al., MLL3/MLL4 are required for CBP/p300 binding on enhancers and super-enhancer formation in brown adipogenesis. Nucleic Acids Res, 2017. 45(11): p. 6388–6403.

13. Lee, J.E., et al., Brd4 binds to active enhancers to control cell identity gene induction in adipogenesis and myogenesis. Nat Commun, 2017. 8(1): p. 2217.

14. Ren, G., et al., Acute depletion of BRG1 reveals its primary function as an activator of transcription. Nat Commun, 2024. 15(1): p. 4561.

15. Lassar, A.B., Finding MyoD and lessons learned along the way. Semin Cell Dev Biol, 2017. 72: p. 3–9.

16. Sartorelli, V. and P.L. Puri, Shaping Gene Expression by Landscaping Chromatin Architecture: Lessons from a Master. Mol Cell, 2018. 71(3): p. 375–388.

17. Sartorelli, V., et al., Molecular mechanisms of myogenic coactivation by p300: direct interaction with the activation domain of MyoD and with the MADS box of MEF2C. Mol Cell Biol, 1997. 17(2): p. 1010–26.

18. de la Serna, I.L., K.A. Carlson, and A.N. Imbalzano, Mammalian SWI/SNF complexes promote MyoD-mediated muscle differentiation. Nat Genet, 2001. 27(2): p. 187–90.

19. Forcales, S.V., et al., Signal-dependent incorporation of MyoD-BAF60c into Brg1-based SWI/SNF chromatin-remodelling complex. EMBO J, 2012. 31(2): p. 301–16.

20. Kimura, E., et al., Cell-lineage regulated myogenesis for dystrophin replacement: a novel therapeutic approach for treatment of muscular dystrophy. Hum Mol Genet, 2008. 17(16): p. 2507–17.

21. Yesbolatova, A., et al., The auxin-inducible degron 2 technology provides sharp degradation control in yeast, mammalian cells, and mice. Nat Commun, 2020. 11(1): p. 5701.

22. Jang, Y., et al., H3.3K4M destabilizes enhancer H3K4 methyltransferases MLL3/MLL4 and impairs adipose tissue development. Nucleic Acids Res, 2019. 47(2): p. 607–620.

23. Froimchuk, E., Y. Jang, and K. Ge, Histone H3 lysine 4 methyltransferase KMT2D. Gene, 2017. 627: p. 337–342.

24. Iurlaro, M., et al., Mammalian SWI/SNF continuously restores local accessibility to chromatin. Nat Genet, 2021. 53(3): p. 279–287.

25. Schick, S., et al., Acute BAF perturbation causes immediate changes in chromatin accessibility. Nat Genet, 2021. 53(3): p. 269–278.

26. Lasko, L.M., et al., Discovery of a selective catalytic p300/CBP inhibitor that targets lineage-specific tumours. Nature, 2017. 550(7674): p. 128–132.

27. Vannam, R., et al., Targeted degradation of the enhancer lysine acetyltransferases CBP and p300. Cell Chem Biol, 2021. 28(4): p. 503–514 e12.

28. Barisic, D., et al., Mammalian ISWI and SWI/SNF selectively mediate binding of distinct transcription factors. Nature, 2019. 569(7754): p. 136–140.

29. Brahma, S. and S. Henikoff, The BAF chromatin remodeler synergizes with RNA polymerase II and transcription factors to evict nucleosomes. Nat Genet, 2024. 56(1): p. 100–111.

30. Martin, B.J.E., et al., Global identification of SWI/SNF targets reveals compensation by EP400. Cell, 2023. 186(24): p. 5290–5307 e26.

31. Narita, T., et al., Enhancers are activated by p300/CBP activity-dependent PIC assembly, RNAPII recruitment, and pause release. Mol Cell, 2021. 81(10): p. 2166–2182 e6.

32. Li, J., et al., Single-Molecule Nanoscopy Elucidates RNA Polymerase II Transcription at Single Genes in Live Cells. Cell, 2019. 178(2): p. 491–506 e28.

33. Hogg, S.J., et al., Targeting histone acetylation dynamics and oncogenic transcription by catalytic P300/CBP inhibition. Mol Cell, 2021. 81(10): p. 2183–2200 e13.

34. Weinert, B.T., et al., Time-Resolved Analysis Reveals Rapid Dynamics and Broad Scope of the CBP/p300 Acetylome. Cell, 2018. 174(1): p. 231–244 e12.

35. Sartorelli, V., et al., Acetylation of MyoD directed by PCAF is necessary for the execution of the muscle program. Mol Cell, 1999. 4(5): p. 725–34.

36. Di Padova, M., et al., MyoD acetylation influences temporal patterns of skeletal muscle gene expression. J Biol Chem, 2007. 282(52): p. 37650–9.

37. Xie, G., et al., MLL3/MLL4 methyltransferase activities control early embryonic development and embryonic stem cell differentiation in a lineage-selective manner. Nat Genet, 2023. 55(4): p. 693–705.

38. Cho, Y.W., et al., Histone methylation regulator PTIP is required for PPARgamma and C/EBPalpha expression and adipogenesis. Cell Metab, 2009. 10(1): p. 27–39.

39. Wang, L., et al., Histone H3K27 methyltransferase Ezh2 represses Wnt genes to facilitate adipogenesis. Proc Natl Acad Sci U S A, 2010. 107(16): p. 7317–22.

40. Zang, C., et al., A clustering approach for identification of enriched domains from histone modification ChIP-Seq data. Bioinformatics, 2009. 25(15): p. 1952–8.

41. Heinz, S., et al., Simple combinations of lineage-determining transcription factors prime cis-regulatory elements required for macrophage and B cell identities. Mol Cell, 2010. 38(4): p. 576–89.

